# Intra-strain elicitation and suppression of plant immunity by *Ralstonia solanacearum* type-III effectors in *Nicotiana benthamiana*

**DOI:** 10.1101/780890

**Authors:** Yuying Sang, Wenjia Yu, Haiyan Zhuang, Yali Wei, Lida Derevnina, Gang Yu, Jiamin Luo, Alberto P. Macho

## Abstract

Effector proteins delivered inside plant cells are powerful weapons for bacterial pathogens, but this exposes the pathogen to potential recognition by the plant immune system. Therefore, the effector repertoire of a given pathogen must be balanced for a successful infection. *Ralstonia solanacearum* is an aggressive pathogen with a large repertoire of secreted effectors. One of these effectors, RipE1, is conserved in most *R. solanacearum* strains sequenced to date. In this work, we found that RipE1 triggers immunity in *N. benthamiana*, which requires the immune regulator SGT1, but not EDS1 or NRCs. Interestingly, RipE1-triggered immunity induces the accumulation of salicylic acid (SA) and the overexpression of several genes encoding phenylalanine-ammonia lyases (PALs), suggesting that the unconventional PAL-mediated pathway is responsible for the observed SA biosynthesis. Surprisingly, RipE1 recognition also induces the expression of jasmonic acid (JA)-responsive genes and JA biosynthesis, suggesting that both SA and JA may act cooperatively in response to RipE1. Finally, we found that RipE1 expression leads to the accumulation of glutathione in plant cells, which precedes the activation of immune responses. *R. solanacearum* encodes another effector, RipAY, which is known to inhibit immune responses by degrading cellular glutathione. Accordingly, we show that RipAY inhibits RipE1-triggered immune responses. This work shows a strategy employed by *R. solanacearum* to counteract the perception of its effector proteins by the plant immune system.

## Introduction

*Ralstonia solanacearum* is considered one of the most destructive plant pathogens, and is able to cause disease in more than 250 plant species (Jiang et al., 2017; Mansfield et al., 2012). As a soil-borne bacterial pathogen, *R. solanacearum* enters plants through the roots, reaches the vascular system, and spreads through xylem vessels, colonizing the plant systemically (Mansfield et al., 2012). This is followed by massive bacterial replication and the disruption of the plant vascular system, leading to eventual plant wilting (Digonnet et al., 2012; Turner et al., 2009).

Most bacterial pathogens deliver proteins inside plant cells via a type-III secretion system (T3SS); such proteins are thus called type-III effectors (T3Es) (Galan et al, 2014). T3Es have been reported to mediate the suppression of basal defenses and the manipulation of plant physiological functions to support bacterial proliferation (Macho et al, 2015; Macho, 2016). Resistant plants have evolved intracellular receptors defined by the presence of nucleotide-binding sites (NBS) and leucine-rich repeat domains (LRRs), thus termed NLRs (Cui et al, 2015). Specific NLRs can detect the activities of specific T3Es, leading to the activation of immune responses, which effectively prevent pathogen proliferation (Chiang & Coaker, 2015). The outcome of these responses is named effector-triggered immunity (ETI), and, in certain cases, may cause a hypersensitive response (HR) that involves the collapse of plant cells. Hormone-mediated signaling plays an essential role in plant immunity. Salicylic acid (SA) is considered the most important hormone in plant immunity against biotrophic pathogens (Vlot *et al*., 2009; Burger & Chory, 2019); Jasmonic acid (JA), on the other hand, is considered the main mediator of immune responses against necrotrophic pathogens (Burger & Chory, 2019). In most cases, both hormones are considered as antagonistic, balancing the effects of each other (Burger & Chory, 2019).

In an evolutionary response to ETI, successful pathogens have acquired T3E activities to suppress this phenomenon (Jones & Dangl, 2006), although reports characterizing T3E suppression of ETI remain scarce, particularly among T3Es within the same strain. While the development of additional T3E activities is a powerful virulence strategy, it also exposes the pathogen to further events of effector recognition. Therefore, the benefits and penalties of T3E secretion need to be finely and dynamically balanced in specific hosts, to ensure the appropriate manipulation of plant functions while evading or suppressing ETI. This balance may be particularly important for *R. solanacearum*, which secretes a larger number of T3Es in comparison to other bacterial plant pathogens (*e.g.* the reference GMI1000 strain is able to secrete more than 70 T3Es) (Peeters et al, 2013).

Plants have evolved to recognize immune elicitors from *R. solanacearum* (Wei et al, 2018; Jayaraman et al., 2016). In terms of mechanism of T3E recognition, the most studied case in *R. solanacearum* is RipP2 (also known as PopP2), which is perceived in Arabidopsis by the RRS1-RPS4 NLR pair (Gassmann et al, 1999; Deslandes et al, 2002; Tasset et al, 2010; Williams et al, 2014; Le Roux et al, 2015; Sarris et al, 2015). Additionally, several *R. solanacearum* T3Es were shown to induce cell death in different plant species (Peeters et al, 2013; Clarke et al, 2015), although, in most cases, it is unclear whether these are due to toxic effects caused by effector overexpression or a host immune response. Some *R. solanacearum* T3Es have also been shown to cause a restriction of host range; such is the case for RipAA and RipP1 (also known as AvrA and PopP1, respectively), which are perceived and restrict host range in *Nicotiana* species (Poueymiro et al, 2009). RipP1 also triggers resistance in petunia (Lavie et al, 2002). Similarly, RipB-triggered immunity has been reported as the major cause for avirulence of *R. solanacearum* RS1000 in *Nicotiana* species (Nakano & Mukaihara, 2019), RipAX2 (also known as Rip36) have been shown to induce resistance in eggplant and its wild relative *Solanum torvum* (Nahar et al, 2014; Morel et al, 2018a), and several T3Es from the AWR family (also known as RipA) restrict bacterial growth in Arabidopsis (Sole et al, 2012). Although the utilization of these recognition systems to generate disease-resistant crops is tantalizing, it is imperative to understand the mechanisms underlying the activation of plant immunity and their potential suppression by other T3Es within *R. solanacearum*.

The *ripE1* gene encodes a protein secreted by the type-III secretion system in the *R. solanacearum* GMI1000 strain (phylotype I) (Mukaihara et al, 2010), and is conserved across *R. solanacearum* strains from different phylotypes (Peeters et al, 2013). Based on sequence analysis, RipE1 is homologous to other T3Es in *Pseudomonas syringae* (HopX) and *Xanthomonas spp* (XopE) (Figure S1; Peeters et al, 2013), belonging to the HopX/AvrPphB T3E family (Nimchuk et al, 2007). This family is characterized by the presence of a putative catalytic triad consisting of specific cysteine, histidine, and aspartic acid residues, which are conserved in RipE1 (Nimchuk et al, 2007; Figure S1), and is similar to several enzyme families from the transglutaminase protein superfamily, such as peptide N-glycanases, phytochelatin synthases, and cysteine proteases (Makarova et al, 1999). AvrPphB, from *P. syringae* pv. *phaseolicola*, the original member of the HopX/AvrPphB family, was identified based on its ability to activate immunity in certain bean cultivars (Mansfield et al, 1994). Divergent members from this family in other strains also trigger immunity, and this requires the putative catalytic cysteine (Nimchuk et al, 2007). Previous sequence analysis of T3Es from the HopX family also identified a conserved domain (domain A) required for HopX induction of immunity in bean and Arabidopsis, which as hypothesized to represent a host-target interaction domain or a novel nucleotide/cofactor binding domain (Nimchuk et al, 2007).

In this work, we studied the impact of RipE1 in plant cells, and found that RipE1 is recognized by the plant immune system in both *N. benthamiana* and Arabidopsis, leading to the activation of immune responses. We further investigate the immune components and signaling pathways associated to this effector recognition. Finally, we found that another effector in *R. solanacearum* GMI1000 is able to inhibit RipE1-triggered immune responses in *N. benthamiana*, explaining the fact that RipE1 does not seem to be an avirulence determinant in this plant species.

## Results

### RipE1 triggers cell death upon transient expression in *Nicotiana benthamiana*

In order to understand the impact of RipE1 in plant cells, we first used an *Agrobacterium tumefaciens* (hereafter, Agrobacterium)-mediated expression system in *Nicotiana benthamiana* leaves to transiently express RipE1 that is fused to a carboxyl-terminal green fluorescent protein (GFP) tag (RipE1-GFP). Two days after Agrobacterium infiltration, we noticed the collapse of infiltrated tissues expressing RipE1-GFP, but not a GFP control (Figure 1a). This tissue collapse correlated with a release of ions from plant cells (Figure 1b), and cell death was confirmed by trypan blue staining (Figure S2). Mutation of the catalytic cysteine to an alanine residue has been shown to disrupt the catalytic activity of enzymes with a catalytic triad similar to that conserved in RipE1 (Gimenez-Ibanez et al, 2014; Figure 1c). To determine if the putative catalytic activity is required for RipE1 induction of cell death, we generated an equivalent mutant in RipE1 (C172A; Figure 1c). We also generated an independent mutant with a deletion on the eight amino acids that constitute the conserved domain A (Nimchuk et al, 2007; Figure 1c). These mutations did not affect the accumulation of RipE1 (Figure 1d), but abolished the induction of tissue collapse and the ion leakage caused by RipE1 expression (Figure 1e and 1f), indicating that RipE1 requires both the catalytic cysteine and the conserved domain A for the induction of cell death in plants.

**Figure 1.**
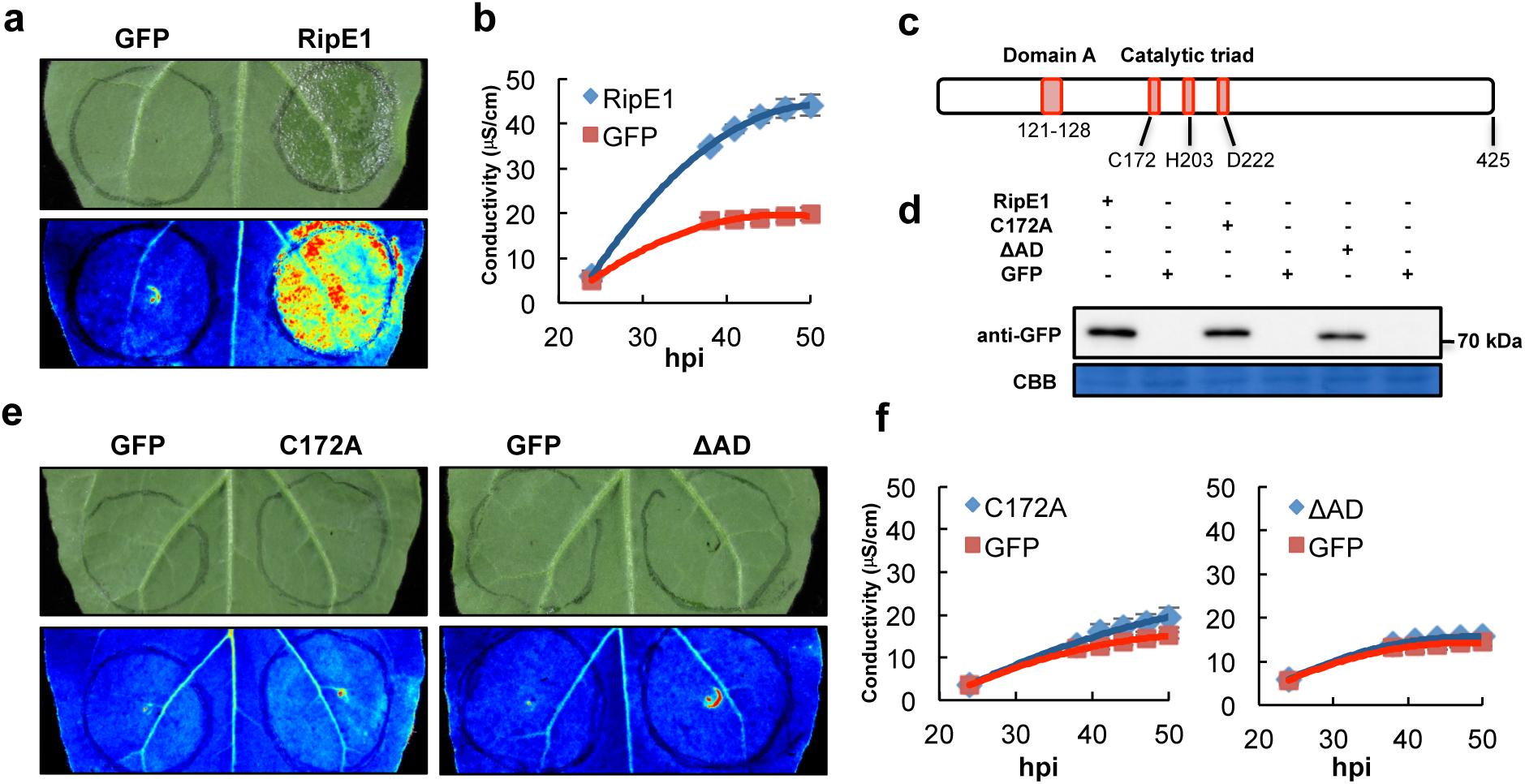
RipE1 triggers cell death in *Nicotiana benthamiana*. (a) RipE1-GFP or GFP (as control) were expressed in the same leaf of *N. benthamiana* using Agrobacterium with an OD_600_ of 0.5. Photos were taken 2 days post-inoculation with a CCD camera (upper panel) or an UV camera (lower panel). UV signal corresponds to the development of cell death (not GFP fluorescence). UV images were taken from the abaxial side and flipped horizontally for representation. (b) Ion leakage measured in leaf discs taken from *N. benthamiana* tissues expressing RipE1-GFP or GFP (as control), representative of cell death, at the indicated time points. Values indicate mean ± SE (n=3 biological replicates). (c) Simplified diagram of RipE1, including the residues comprising the Domain A and the predicted catalytic triad. (d) Western blot showing the accumulation of RipE1 mutant variants. ΔAD corresponds to a deletion mutant of the Domain A (residues 121-128). Molecular weight (kDa) marker bands are indicated for reference. (e) Cell death triggered by RipE1 mutant variants (conditions as in (a)). (f) Ion leakage measured in leaf discs taken from *N. benthamiana* tissues expressing RipE1 mutant variants (conditions as in (b). Each experiment was repeated at least 3 times with similar results.

Interestingly, RipE1 was also identified in a systematic screen performed in our laboratory to identify *R. solanacearum* T3Es that suppress immune responses triggered by bacterial elicitors. In this screen we found that RipE1 expression suppresses the burst of reactive oxygen species (ROS) and the activation of mitogen-activated protein kinases (MAPKs) triggered upon treatment with the bacterial flagellin epitope flg22, which acts as an immune elicitor (Figure S3a and S3b). RipE1 requires both the catalytic cysteine and the conserved domain A for this activity (Figure S3c). However, we considered the possibility that these responses are abolished by the death of plant cells rather than an active immune suppression. Time-course experiments showed that the suppression of flg22-triggered ROS correlated with the appearance of cell death (Figure S3a and S3d), making it difficult to uncouple these observations.

### RipE1 activates salicylic acid-dependent immunity in *N. benthamiana*

The induction of cell death by pathogen effectors may reflect toxicity in plant cells or the activation of immune responses that lead to a HR. Salicylic Acid (SA) plays a major role in the activation of immune responses after the perception of different types of invasion patterns (Vlot *et al*., 2009). To determine whether RipE1 activates immune responses, we first measured the expression of the *N. benthamiana* ortholog of the Arabidopsis gene *PATHOGENESIS-RELATED-1* (*PR1*), which is a hallmark of SA-dependent immune responses (Vlot et al., 2009, Ward *et al*., 1991). Expression of RipE1-GFP (but not the C172A catalytic mutant) significantly enhanced the accumulation of *NbPR1* transcripts (Figure 2a). In keeping with the notion that RipE1 activates a defense response against *R. solanacearum*, RipE1 expression in *N. benthamiana* leaves enhanced resistance against subsequently inoculated *R. solanacearum* Y45, which is otherwise pathogenic in *N. benthamiana* (Li *et al*., 2011) (Figure 2b). The bacterial salicylate hydroxylase NahG converts SA to catechol, which leads to the suppression of SA-dependent responses (Delaney et al., 1994). The expression of NahG-GFP in *N. benthamiana* slightly enhanced the accumulation of RipE1 fused to a carboxyl-terminal N-luciferase tag (Nluc) (Figure S4), consistent with the reported role of SA in hindering Agrobacterium-mediated transformation (Rosas-Diaz et al, 2016); despite this, *NahG* expression partially suppressed RipE1-triggered cell death, ion leakage, and *NbPR1* expression (Figure 2c, d and e). Altogether, these data suggest that RipE1 induces SA-dependent immune responses in plant cells, which cause the development of a HR.

**Figure 2.**
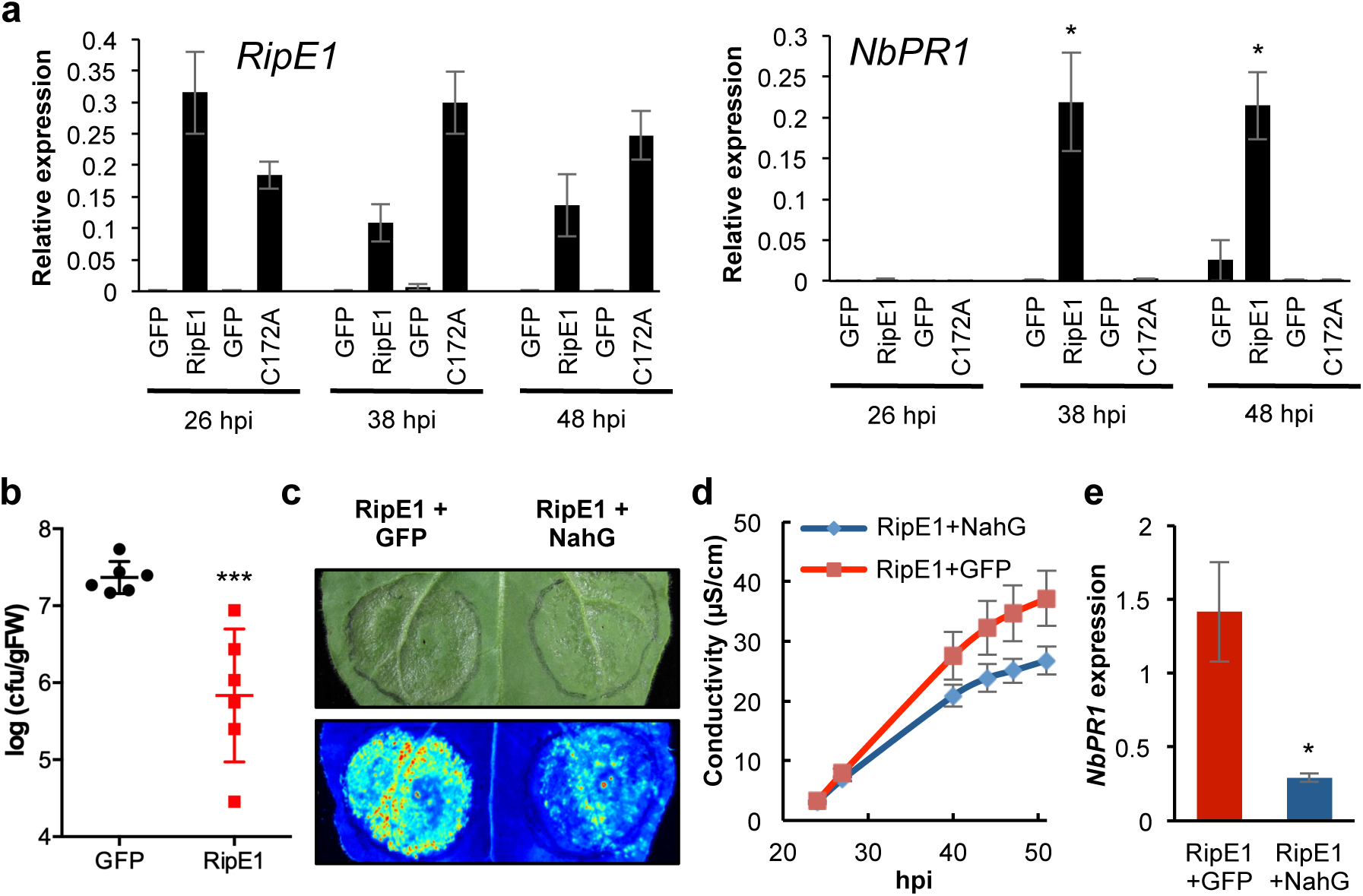
RipE1 activates SA-dependent immune responses in *N. benthamiana*. (a) Quantitative RT-PCR to determine the expression of *RipE1* and *NbPR1* in *N. benthamiana* tissues expressing GFP, RipE1, or RipE1 C172A, using Agrobacterium with an OD_600_ of 0.1. Samples were taken at the indicated times after Agrobacterium infiltration. In each case, the RipE1 variants and their respective GFP control were expressed in the same leaf, and values are represented side-by-side. Expression values are relative to the expression of the housekeeping gene *NbEF1a*. Values indicate mean ± SE (n=3 biological replicates). (b) RipE1-GFP or GFP (as control) were expressed in the same leaf of *N. benthamiana* using Agrobacterium with an OD_600_ of 0.5. Twenty-four hours after Agrobacterium infiltration, before the appearance of cell death, a 10^5^ cfu/ml inoculum of *R. solanacearum* Y45 was infiltrated into the same tissues. Samples were taken one day post-inoculation to determine Y45 colony-forming units (cfu) per gram of tissue. Values indicate mean ± SE (n=6 biological replicates). (c-e) RipE1-Nluc was co-expressed with GFP (as control) or with NahG-GFP in the same leaf. Protein accumulation is shown in the Figure S4. (c) Photos were taken 2.5 days post-inoculation with a CCD camera (upper panel) or an UV camera (lower panel). UV signal corresponds to the development of cell death (not GFP fluorescence). UV images were taken from the abaxial side and flipped horizontally for representation. (d) Ion leakage measured in leaf discs taken from *N. benthamiana* tissues expressing RipE1 together with GFP or NahG-GFP, representative of cell death, at the indicated time points. Values indicate mean ± SE (n=3 biological replicates). (e) Quantitative RT-PCR to determine the expression of *NbPR1* in *N. benthamiana* tissues 48 hours after Agrobacterium infiltration. Expression values are relative to the expression of the housekeeping gene *NbEF1a*. Values indicate mean ± SE (n=3 biological replicates). Asterisks indicate significant differences compared to the mock control according to a Student’s t test (* p < 0.05; *** p < 0.001). Each experiment was repeated at least 3 times with similar results.

### RipE1 enhances the expression of *PAL* genes and the biosynthesis of salicylic acid and jasmonic acid

The expression of RipE1 led to a dramatic increase in SA accumulation in *N. benthamiana* (Figure 3a), consistent with the observed overexpression of *NbPR1* (Figure 2a). This reinforces the idea that RipE1 is perceived by the plant immune system and this leads to the activation of SA biosynthesis and SA-dependent immune responses. In Arabidopsis, the chloroplastic pathway mediated by isochorismate synthethase 1 (ICS1) plays a predominant role in the pathogen-induced SA biosynthesis (Wildermuth et al, Nature, 2001; Garcion et al, Plant Physiology, 2008). However, gene expression analysis showed that the expression of the *N. benthamiana* ortholog of the Arabidopsis *ICS1*, *NbICS1*, was significantly reduced upon RipE1 expression (Figure 3b), despite the simultaneous high *NbPR1* transcript accumulation (Figure 2a). SA can also be synthesized from phenylalanine in a pathway mediated by phenylalanine ammonia lyases (PALs). In contrast with the expression of *NbICS1*, several genes encoding NbPALs were up-regulated upon expression of RipE1, but not the catalytic mutant version (Figure 3c-e), suggesting that this pathway may mediate the enhancement of SA biosynthesis upon perception of RipE1 activity. SA and Jasmonic Acid (JA) are considered antagonistic hormones in plant immune responses. Surprisingly, instead of a reduction of JA-associated gene expression, we found a slight increase in the accumulation of *NbLOX2* transcripts in early time points upon expression of catalytically active RipE1 (Figure 3f). In Arabidopsis, LOX2 contributes to the biosynthesis of JA (Bell et al, 1995). Accordingly, we detected an increase in JA contents upon RipE1 expression (Figure S5), indicating that RipE1 perception does not inhibit JA signalling, but rather leads to an enhancement of JA biosynthesis and associated gene expression.

**Figure 3.**
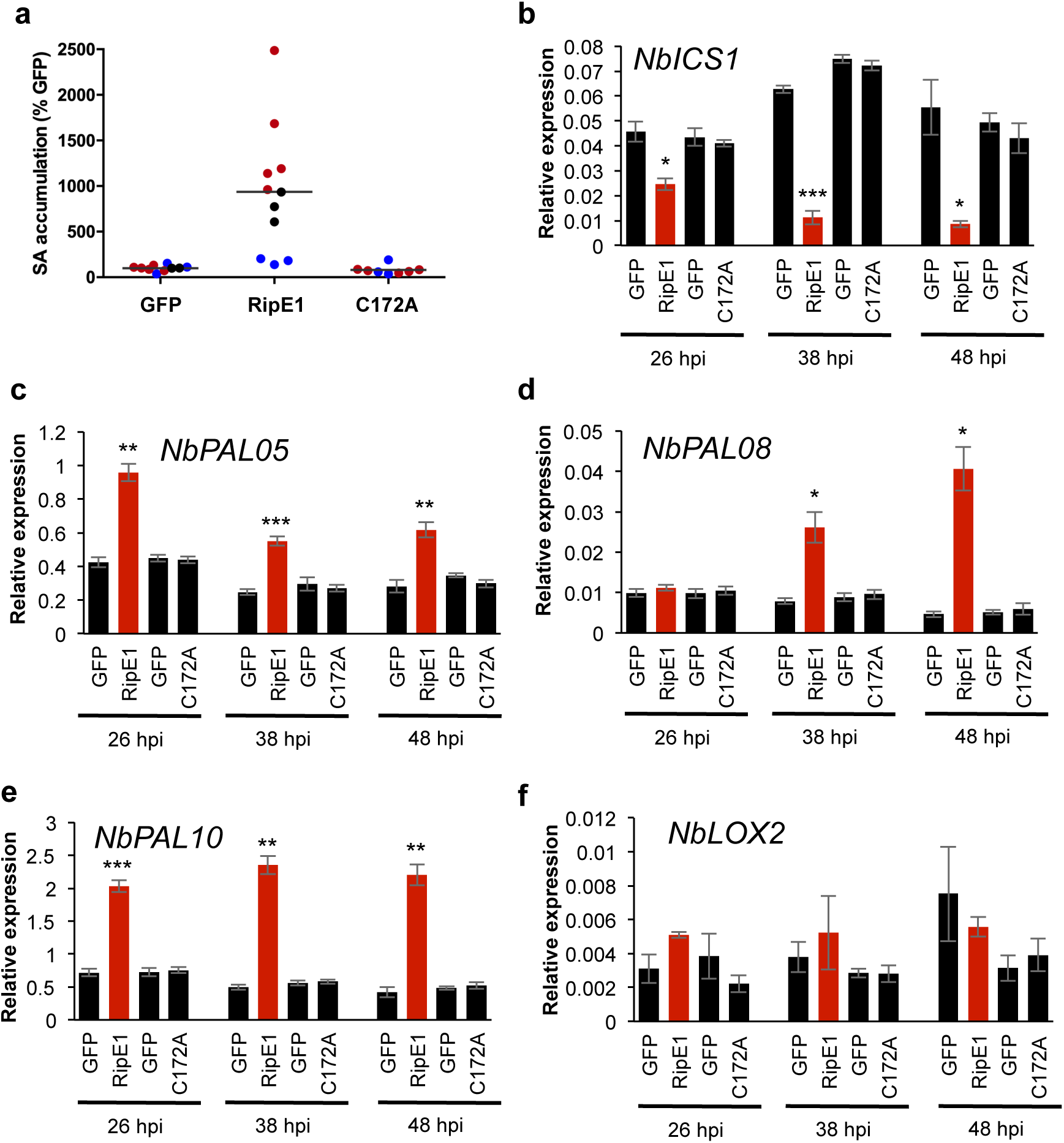
RipE1 perception enhances the expression of *PAL* genes and SA biosynthesis in *N. benthamiana*. (a) Measurement of SA accumulation in *N. benthamiana* tissues expressing GFP, RipE1, or RipE1 C172A, using Agrobacterium with an OD_600_ of 0.5. Samples were taken 42 hours after Agrobacterium infiltration. Three independent biological repeats were performed, and the different colors indicate values from different replicates. Values are represented as % of the GFP control in each replicate. (b-f) Quantitative RT-PCR to determine the expression of *NbICS1* (b), *NbPAL05* (c), *NbPAL08* (d), *NbPAL10* (e), and *NbLOX2* (f) in *N. benthamiana* tissues expressing GFP, RipE1, or RipE1 C172A, using Agrobacterium with an OD_600_ of 0.5. Samples were taken at the indicated times after Agrobacterium infiltration. In each case, the RipE1 variants and their respective GFP control were expressed in the same leaf, and values are represented side-by-side. *NbPAL05:* Niben101Scf05617g00005.1; *NbPAL08*: Niben101Scf03712g01008.1; *NbPAL10:* Niben101Scf12881g00010.1. Expression values are relative to the expression of the housekeeping gene *NbEF1a*. Values indicate mean ± SE (n=3 biological replicates). Asterisks indicate significant differences compared to the mock control according to a Student’s t test (* p < 0.05; ** p < 0.01; *** p < 0.001). Each experiment was repeated at least 3 times with similar results.

### RipE1-triggered immunity requires SGT1, but not EDS1 or NRC proteins

The suppressor of the G2 allele of *skp1* (SGT1) plays an essential role in ETI, and is required for the induction of disease resistance mediated by most NLRs (Azevedo *et al*., 2002; Kadota *et al*., 2010). Virus-induced gene silencing (VIGS) of *NbSGT1* abolished RipE1-triggered cell death, ion leakage, and *NbPR1* expression (Figure 4a-d), indicating that RipE1-triggered immunity requires SGT1. While most NLRs require SGT1 to function, a specific group of NLRs containing an N-terminal Toll-like interleukin-1 receptor (TIR) domain also require EDS1 (Wiermer et al, 2005; Schultink et al, 2017). *N. benthamiana* plants carrying a stable knockout mutation in *EDS1* (Schultink et al, 2017) displayed clear RipE1-triggered cell death (Figure 4e), suggesting that RipE1-triggered immunity is not mediated by a TIR-NLR. Other NLRs contain a C-terminal coiled coil (CC) domain, and a specific subset of CC-NLRs require a network of helper NLRs termed NRC proteins (Wu et al, 2016). Interestingly, silencing of NRC proteins did not impact RipE1-triggered cell death (Figure S6), suggesting that RipE1-triggered immunity is not mediated by an NLR within the NRC network.

**Figure 4.**
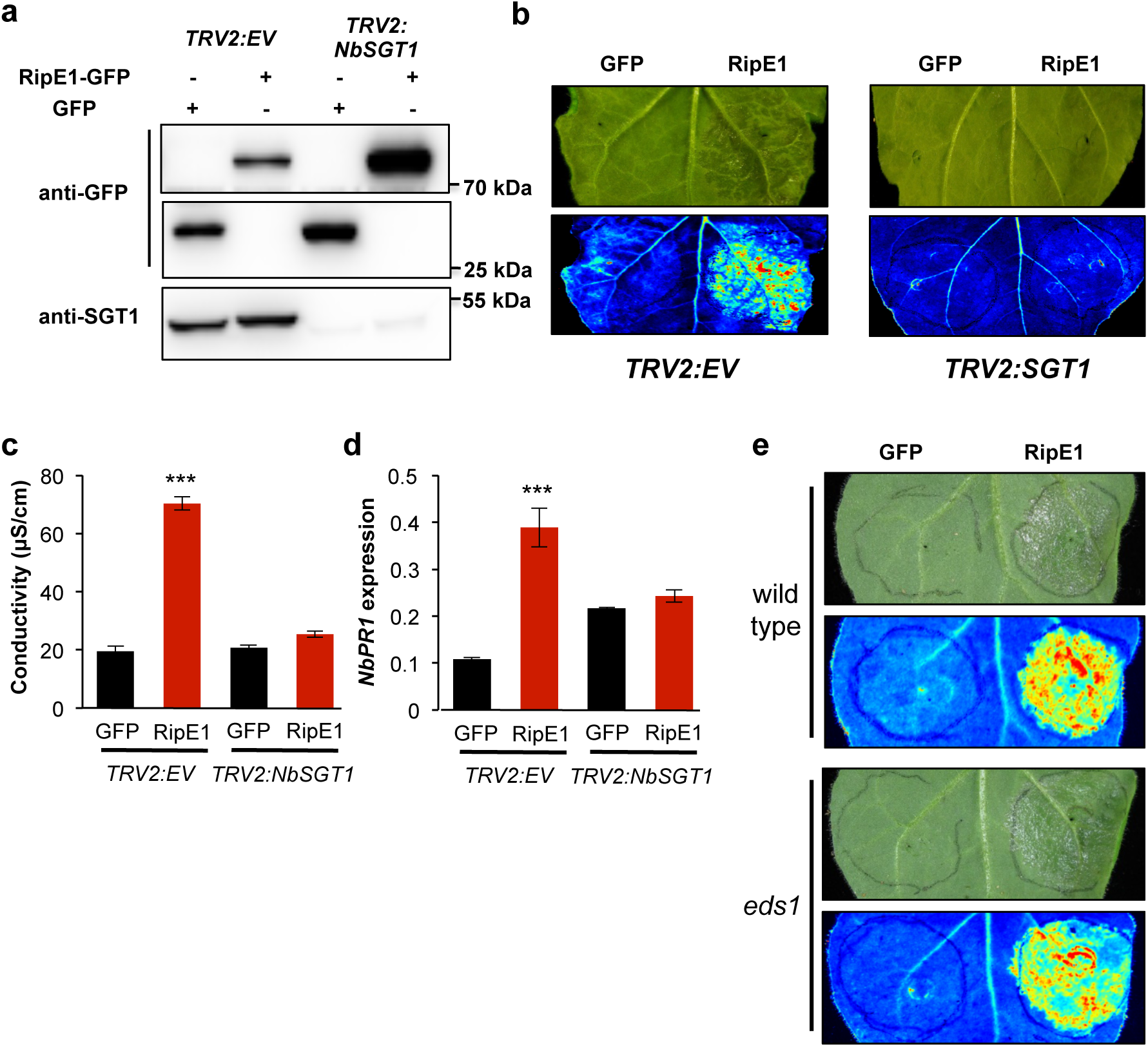
RipE1-triggered responses require SGT1, but not EDS1. (a-d) RipE1-GFP or GFP (as control) were expressed in the same leaf of *N. benthamiana* undergoing VIGS of *NbSGT1* or VIGS with an empty vector (EV) construct (as control), using Agrobacterium with an OD_600_ of 0.5. (a) Western blot showing the accumulation of GFP, RipE1-GFP, and endogenous *NbSGT1*. Molecular weight (kDa) marker bands are indicated for reference. (b) Photos were taken 2 days post-inoculation with a CCD camera (upper panel) or an UV camera (lower panel). UV signal corresponds to the development of cell death (not GFP fluorescence). UV images were taken from the abaxial side and flipped horizontally for representation. (c) Ion leakage measured in leaf discs taken from *N. benthamiana* tissues expressing RipE1-GFP or GFP (as control), representative of cell death, 48 hours after Agrobacterium infiltration. Values indicate mean ± SE (n=3 biological replicates). (d) Quantitative RT-PCR to determine the expression of *NbPR1* in *N. benthamiana* tissues 48 hours after Agrobacterium infiltration. Expression values are relative to the expression of the housekeeping gene *NbEF1a*. Values indicate mean ± SE (n=3 biological replicates). (e) RipE1-GFP or GFP (as control) were expressed in the same leaf of *N. benthamiana* wild type or a stable *eds1* knockout mutant, using Agrobacterium with an OD_600_ of 0.5. Photos were taken 2 days post-inoculation with a CCD camera (upper panel) or an UV camera (lower panel). UV signal corresponds to the development of cell death (not GFP fluorescence). UV images were taken from the abaxial side and flipped horizontally for representation. Asterisks indicate significant differences compared to the mock control according to a Student’s t test (*** p < 0.001). Each experiment was repeated at least 3 times with similar results.

### RipE1 activates immunity in Arabidopsis

Arabidopsis transgenic plants expressing RipE1-GFP from a *35S* inducible promoter died after germination (data not shown). Therefore, we generated Arabidopsis transgenic plants expressing RipE1-GFP and RipE1^C172A^-GFP from an estradiol (EST)-inducible promoter. Five-week-old plants expressing RipE1-GFP, but not RipE1^C172A^-GFP, showed reduced growth in soil upon EST treatment for 14 days (Figure 5a). To determine whether RipE1-triggered growth reduction in Arabidopsis correlates with the activation of immunity, we first monitored the expression of defence-related genes. Similar to the result observed upon expression in *N. benthamiana*, expression of RipE1 in Arabidopsis triggered the overexpression of *AtPR1* (Figure 5b). However, in Arabidopsis, the enhanced *PR1* expression correlated with an overexpression of *AtICS1*, but not *AtPAL1*, upon RipE1 expression (Figure 5b). As observed in *N. benthamiana*, RipE1 expression led to the overexpression of the JA marker genes *AtVSP2* and *AtPDF1*.2 (Figure 5b). This indicates that, as observed in *N. benthamiana*, RipE1 activates SA- and JA-dependent signalling in Arabidopsis. To determine whether the activation of defence-related genes in Arabidopsis leads to an efficient immune response against *R. solanacearum*, we inoculated RipE1-expressing plants by soil-drenching with *R. solanacearum* after EST treatment for 2 days. As shown in the figure 5c, RipE1-expressing plants displayed weaker and delayed disease symptoms upon *R. solanacearum* inoculation, reflecting an enhanced disease resistance upon *RipE1* expression. RipE1-expressing plants also showed a moderate reduction in bacterial growth after *R. solanacearum* infiltration in the leaves (Figure S7a), suggesting that the immune response is not exclusively associated to invasion or proliferation in the root. However, RipE1-expressing plants did not display enhanced resistance against the leaf-borne pathogen *Pseudomonas syringae* pv. *tomato* DC3000 (Figure S7b and S7c).

**Figure 5.**
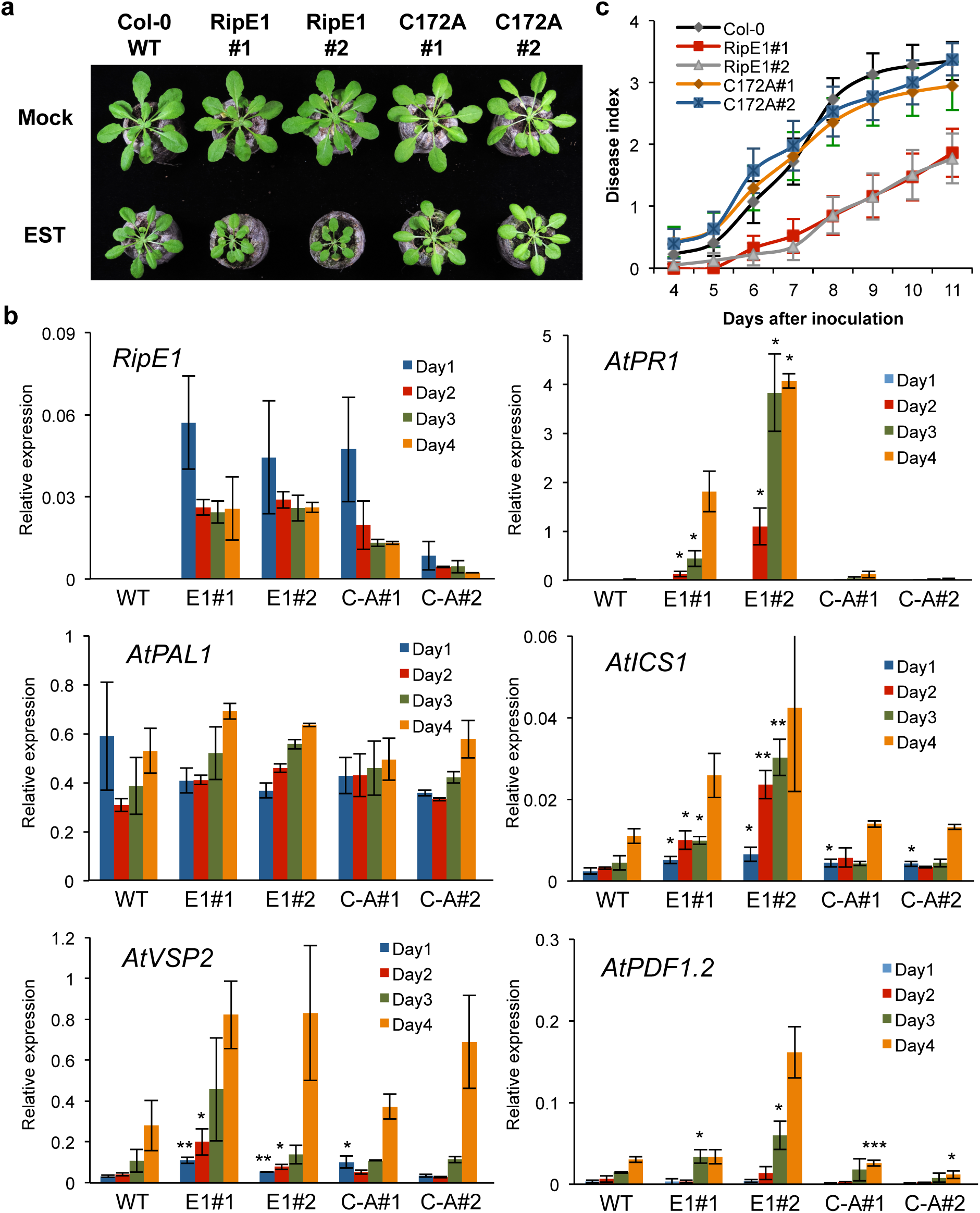
RipE1 triggers immunity in Arabidopsis. (a) Arabidopsis Col-0 wild type or independent stable transgenic lines expressing RipE1 or RipE1 C172A from an estradiol (EST)-inducible promoter were grown for 3 weeks and then treated sprayed with 100 µM EST daily. Photographs were taken 2 weeks after beginning the EST treatment. (b) Arabidopsis 4 day-old seedlings were treated with 25 µM EST and samples were taken 1, 2, 3, or 4 days after EST treatment. Quantitative RT-PCR to determine the expression of *RipE1, AtPR1, AtPAL1, AtICS1, AtVSP2,* and *AtPDF1.2*. Expression values are relative to the expression of the housekeeping gene *AtACT2*. Values indicate mean ± SE (n=3 biological replicates). (c) Arabidopsis Col-0 wild type or EST-RipE1 transgenic plants were grown for 4 weeks and then treated with 100 µM EST for 2 days before inoculation with *R. solanacearum* GMI1000 by soil-drenching. Plants showed no difference in root or shoot size at the time of inoculation. The results are represented as disease progression, showing the average wilting symptoms in a scale from 0 to 4 (mean ± SEM). n=20 plants per genotype. Asterisks indicate significant differences compared to the mock control according to a Student’s t test (* p < 0.05; ** p < 0.01; *** p < 0.001). Each experiment was repeated at least 3 times with similar results.

### RipE1-triggered immune responses are suppressed by RipAY

RipE1 expression activates immunity in Arabidopsis and *N. benthamiana*, although both plant species are susceptible hosts for *R. solanacearum* GMI1000 (or a derivative strain carrying mutations in *ripP1* and *ripAA*, in the case of *N. benthamiana*; Poueymiro et al, 2009), which carries RipE1. Therefore, we reasoned that other T3E(s) in GMI1000 may be able to suppress RipE1-triggered immunity in the context of infection. We recently identified a *R. solanacearum* T3E, RipAY, which is able to suppress SA-dependent immune responses through the degradation of glutathione (Sang et al, 2016; Mukaihara et al, 2016); however, the ability of RipAY to suppress immunity triggered by other *R. solanacearum* T3Es remained unknown. Interestingly, the expression of RipE1 in *N. benthamiana* leads to an increase in glutathione accumulation in plant tissues, which precedes the onset of immune responses (Figure 6a). Considering that both RipAY and RipE1 are present in GMI1000, we sought to determine if RipAY has the ability to suppress RipE1-triggered immunity. Indeed, expression of RipAY in *N. benthamiana* did not affect the accumulation of RipE1 (Figure S8), but abolished the tissue collapse and ion leakage caused by *RipE1* expression (Figure 6b and c). Moreover, RipAY was able to suppress the overexpression of several SA-related genes triggered by RipE1 (Figure 6d and S9), indicating that RipAY suppresses RipE1-triggered immune responses. RipAY did not significantly suppress the RipE1-triggered expression of *NbLOX2* (Figure S9), although this may be due to the weaker induction of this gene compared to the tested SA-related genes (Figure 3 and S9). Interestingly, however, a RipAY point mutant unable to degrade glutathione (RipAY^E216Q^; Sang et al, 2016) did not suppress RipE1-triggered responses (Figure 6b-d), suggesting that RipAY suppresses RipE1-triggered immunity through the degradation of cellular glutathione.

**Figure 6.**
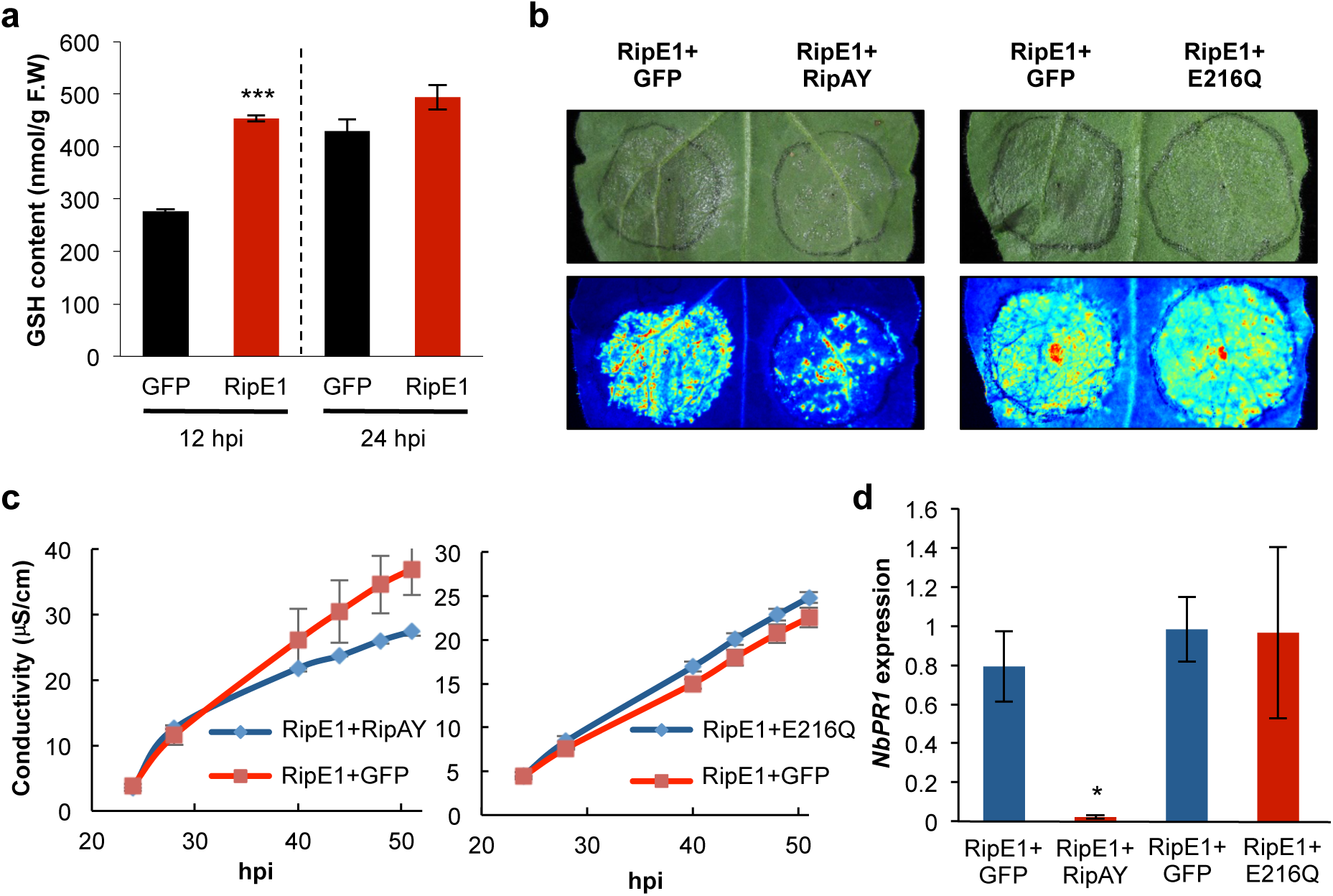
RipE1-triggered immune responses are suppressed by RipAY. (a) RipE1-GFP or GFP (as control) were expressed in the same leaf of *N. benthamiana* using Agrobacterium with an OD_600_ of 0.5, and samples were taken at the indicated time points to measure the accumulation of glutathione (GSH). (b-d) RipE1-Nluc was co-expressed with GFP (as control), with RipAY-GFP, or with RipAY-E216Q-GFP, respectively, in the same leaf. Protein accumulation is shown in the Figure S8. (b) Photos were taken 2.5 days post-inoculation with a CCD camera (upper panel) or an UV camera (lower panel). UV signal corresponds to the development of cell death (not GFP fluorescence). UV images were taken from the abaxial side and flipped horizontally for representation. (c) Ion leakage measured in leaf discs taken from *N. benthamiana* tissues expressing RipE1 together with GFP or RipAY-GFP, representative of cell death, at the indicated time points. Values indicate mean ± SE (n=3 biological replicates). (d) Quantitative RT-PCR to determine the expression of *NbPR1* in *N. benthamiana* tissues 48 hours after Agrobacterium infiltration. Expression values are relative to the expression of the housekeeping gene *NbEF1a*. Values indicate mean ± SE (n=3 biological replicates). Asterisks indicate significant differences compared to the mock control according to a Student’s t test (* p < 0.05; *** p < 0.001). Each experiment was repeated at least 3 times with similar results.

## Discussion

Expression of T3Es in plant cells may either induce cell death because of cell toxicity or lead to the activation of an immunity-associated HR. Over-expression of RipE1 in *N. benthamiana* leads to a HR that: (*i*) is dependent on the immune regulator SGT1; (*ii*) activates SA accumulation and *PR1* expression; (*iii*) restricts growth of *R. solanacearum* Y45; and (*iv*) is suppressed by the NahG and other *R. solanacearum* effectors, indicating that RipE1-mediated cell death is due to the activation of immunity in the host. It is, however, noteworthy that cell death induced by RipE1 develops slower than that triggered by other HR-inducing T3Es (*i.e.* RipAA; Figure S2). Several T3Es within the HopX/AvrPphB family are predicted enzymes that are associated with activation of host immunity, although the association of the predicted catalytic activity with the activation of immunity seems to be differ among them. While the ability of AvrPphB and several other family members to trigger immunity requires the putative catalytic cysteine (Mansfield et al, 1994; Nimchuk et al, 2007), other members with the predicted catalytic activity, such as HopX from *P. syringae* pv *tabaci* or *P. syringae* pv *phaseolicola* race 6, do not trigger immunity in the same hosts (Stevens et al, 1998; Nimchuk et al, 2007). In the case of RipE1, the putative catalytic cysteine is required for the induction of immunity, which suggests that RipE1 is an active enzyme, and that this catalytic activity leads to perception by the host immune system. Moreover, the conserved domain A (Nimchuk et al, 2007) is also required for the activation of immunity by RipE1. In addition, we found that RipE1 is able to suppress elicitor-triggered immune responses in *N. benthamiana*. However, since this activity correlates with the induction of cell death, it is difficult to uncouple both observations, and further studies on the virulence activity of RipE1 will require the utilization of a host plant that is unable to recognize it.

The fact that RipE1 is recognized, and activate immune responses, in both *N. benthamiana* and Arabidopsis suggests at least two scenarios: it is possible that the NLR responsible for this recognition is conserved in both species; on the other hand, it is also possible that both species have independently develop NLRs that recognize RipE1. Although we did not identify the NLR involved, we determined that, at least in *N. benthamiana*, RipE1 recognition does not rely on EDS1 or the NRC network, pointing to a CC- NRC-independent NLR. Interestingly, although RipE1 perception leads to the accumulation of SA in both plant species, the associated gene expression patterns seem to differ. The ICS pathway plays a predominant role in the pathogen-induced SA biosynthesis in Arabidopsis (Wildermuth et al, Nature, 2001; Garcion et al, Plant Physiology, 2008). In agreement with this, the RipE1-triggered overexpression of *AtPR1* in Arabidopsis correlates with an enhanced expression of *AtICS1*, but not *AtPAL1*. However, it seems that the RipE1-induced increase in SA content in *N. benthamiana* correlates with a reduction of *NbICS1* gene expression, and an increase in the expression of several *NbPAL* genes. Considering that ICS1 is normally regulated at the transcriptional level upon pathogen perception (Wildemurth et al, 2001), our results suggest that the PAL pathway is more relevant than the ICS pathway for the induction of RipE1-triggered immunity in *N. benthamiana*, indicating that both pathways are differentially required for distinct immune responses in different plant species. Similarly, both the ICS and PAL pathways have been reported to be required for pathogen-induced SA biosynthesis in soybean (Shine et al, 2016). The reduction in ICS1 expression in *N. benthamiana* may reflect a compensatory effect between the ICS and PAL pathway. In addition to different gene expression patterns, the physiological output in both plant species may be different. Although RipE1 expression caused an inhibition of Arabidopsis growth, we did not observe any signs of cell death (data not shown), which contrasts with our observation in *N. benthamiana*. However, this may be caused by differences in the expression system used in both plants (Agrobacterium-mediated transient expression in *N. benthamiana vs* EST-induced expression in Arabidopsis stable transgenic plants).

Another surprising aspect of RipE1-triggered immunity is the fact that it leads to the simultaneous accumulation of SA and JA, and to a strong and moderate SA- and JA-triggered gene expression, respectively, in both *N. benthamiana* and Arabidopsis. This suggests that, in the case of RipE1-triggered immunity, SA and JA may play a cooperative role, possibly reflecting the complexity of the *R. solanacearum* infection process compared to other pathogens. In keeping with this notion, although RipE1-expressing Arabidopsis plants displayed enhanced resistance against *R. solanacearum* and up-regulation of SA-related genes, they did not show enhanced resistance against the leaf-borne pathogen *P. syringae pv. tomato* DC3000 (Figure S6). Since the enhancement of JA signalling has been associated to a promotion of virulence by this pathogen (Gimenez-Ibanez *et al*, 2016), the observed up-regulation of JA-related genes may underlie this phenomenon.

If RipE1 triggers immunity in *N. benthamiana*, why is it that a GMI1000 strain without RipP1 and RipAA (but having RipE1) can cause a successful infection in *N. benthamiana* without triggering immunity (Poueymiro et al, 2009)? Here, we found that another effector within GMI1000, RipAY, is able to inhibit RipE1-triggered immunity. Since RipE1 perception correlates with an enhancement of cellular glutathione, and RipAY requires its gamma-glutamyl cyclotransferase activity to inhibit RipE1-triggered HR, the degradation of glutathione or other gamma-glutamyl compounds (Sang et al, 2016; Mukaihara et al, 2016; Fujiwara et al, 2016) is the most likely mechanism for this inhibition. Besides RipAY, other T3Es within GMI1000 likely contribute to the suppression of RipE1-triggered HR by targeting other immune functions (Yu et al, bioRxiv, 2019; Wang & Macho, unpublished data). This reflects bacterial adaptation: RipE1 could be important for virulence, but also triggers immunity. In this context, instead of losing RipE1, *R. solanacearum* has developed other effectors to suppress the induction of immunity, while keeping RipE1 virulence activity. This is reminiscent of what has been shown for *P. syringae* pv. *syringae* B728a, where several effectors within the same strain suppress the HR triggered by HopZ3, which otherwise acts as a virulence factor (Rufian et al, 2018). Similarly, although transient expression of HopX from *P. syringae* pv *tomato* (*Pto*) triggers HR in specific Arabidopsis accessions, it does not trigger HR in the context of *Pto* infection (Nimchuk et al, 2007). It is possible that, as in the case of RipE1, the immune responses triggered by HopX are masked during *Pto* infection (as suggested in Nimchuk et al, 2007), likely due to the suppression by other effectors within the same strain.

## Materials and Methods

### Plant materials and growth conditions

*N. benthamiana* plants were grown on soil at one plant per pot in an environmentally controlled growth room at 25 °C under a 16-h light/8-h dark photoperiod with a light-intensity of 130 mE m^-2^s^-1^. *A. thaliana* plants were grown under the same conditions as *N. benthamiana* for collection of seeds. For bacterial virulence and ROS burst assays, *A. thaliana* plants were grown in a growth chamber controlled at 22°C with a 10 h photoperiod and a light-intensity of 100-150 mE m^-2^s^-1^. After *R. solanacearum* inoculation, Arabidopsis plants were transferred to a growth chamber at 27°C with 75% humid under a 12-h light/12-h dark photoperiod.

### Chemicals

The flg22 peptide (TRLSSGLKINSAKDDAAGLQIA) was purchased from Abclonal, USA. All other chemicals were purchased from Sigma-Aldrich unless otherwise stated.

### Plasmids, bacterial strains and cultivation conditions

*R. solanacearum* GMI1000 was grown on solid BG medium plates or cultivated over-night in liquid BG medium at 28°C (Morel et al., 2018b). The *ripE1* gene from *R. solanacearum* GMI1000 cloned in pDONR207 (donated by Nemo Peeters and Anne-Claire Cazale) was subcloned into pGWB505 by LR reaction (ThermoFisher, USA) to generate a fusion protein with eGFP tag at the C-terminal (Nakagawa *et al*., 2007). *RipE1* and *ripE1* mutants were inserted between BamHI and XhoI restriction sites on sXVE:GFPc:Bar estradiol inducible vector using enzyme digestion (Schlücking et al., 2013). These generated binary vectors were transformed into *Agrobacterium tumefaciens* (Agrobacterium) GV3101 *f*or transient or stable gene expression in *N. benthamiana* and *A. thaliana* plants. Agrobacterium carrying pGWB505 vectors were grown at 28°C and 220 rpm in LB medium supplemented with rifampicin 50 mg/l, gentamycin 25 mg/l and spectinomycin 50 mg/l, while those carrying estradiol inducible vectors were grown in rifampicin 50 mg/l, gentamycin 25 mg/l and kanamycin 50 mg/l.

### Site-directed mutagenesis

*RipE1_C172A_* and *RipE1 ΔAD* mutant variants were generated using the QuickChange Lightning Site-Directed Mutagenesis Kit (Life technologies, USA) following the manufacturer’s instructions. RipE1/pDONR207 plasmid was used as template. Primers used for the mutagenesis are listed in Table S1.

### Agrobacterium-mediated gene expression in *A. thaliana and N. benthamiana*

Stable transgenic Arabidopsis plants with *RipE1* and *RipE1* mutated variants driven by estradiol inducible promoter were obtained using the floral dip method (Zhang et. al, 2006). Homozygous T_3_ lines were used for all the experiments. Agrobacterium-mediated transient expression in *N. benthamiana* was performed as described (Li, 2011). Agrobacterium carrying the resultant plasmids were suspended in infiltration buffer to a final OD_600_ of 0.1∼0.5 and infiltrated into the abaxial side of the leaves using the 1 ml needless syringe. Leaf samples were taken at 1-3 dpi (days post infiltration) for analysis based on experimental requirements.

### Protein extraction and western blots

Plant tissues were collected into 2 ml tubes with metal beads and frozen in liquid nitrogen. After grinding with a tissue lyser (Qiagen, Germany) for 1 min at 30 rpm/s, proteins were extracted using protein extraction buffer (100 mM Tris-HCl pH 8, 150 mM NaCl, 10% glycerol, 5 mM Ethylene diamine tetra acetic acid (EDTA), 2 mM Dithiothreitol (DTT), 1x Plant Protease Inhibitor cocktail, 1% NP-40, 2 mM Phenylmethylsulfonyl fluoride (PMSF), 10 mM Na_2_MoO_4_, 10 mM NaF, 2 mM Na_3_VO_4_) and incubating for 5 min. After centrifugation, the supernatants were mixed with SDS loading buffer, incubated at 70 °C for 10 min, and resolved using SDS-PAGE. Proteins were transferred to a PVDF membrane and monitored by western blot using anti-GFP (Abicode, M0802-3a) and anti-luciferase (Sigma, L0159) antibodies.

### Measurement of ROS generation and MAPK activation

PAMP-triggered ROS burst and MAPK activation in plant leaves were measured as described previously (Sang et al., 2017; Segonzac *et al*., 2011). ROS was elicited with 50 nM flg22. MAPK activation assays were performed using 4 to 5-week-old *N. benthamiana*. Two days after Agrobacterium infiltration at OD_600_ of 0.1, the intact leaves were elicited for 15 min after vacuum infiltration of 100 nM flg22. Leaf discs were taken to monitor MAPK activation by western blot with Phospho-p44/42 MAPK (Erk1/2; Thr-202/Tyr-204) antibodies.

### Cell death measurement

Cell death in plant leaves was quantified as previously described (Yu et al, bioRxiv, 2019) by measuring the electrolyte leakage using a conductivity meter (ThermoFisher, USA) or observing the autofluorescence using the BioRad Gel Imager (Bio-Rad, USA). Briefly, one day after Agrobacterium infiltration in *N. benthamiana*, one 13 mm leaf disk was immersed in 4 ml of distilled water for 1 h with gentle shaking and then transferred to a 6-well culture plate containing 4 ml distilled water in each well. The ion conductivity was then measured at different time intervals. Autofluorescence in intact *N. benthamiana* leaves was measured at 2.5 dpi. Trypan blue staining was performed as previously described (Lv *et al*, 2019).

### RNA isolation and qRT-PCR

Five-to-eight day-old Arabidopsis seedlings were grown in sterile conditions and 8-10 seedlings grown in an independent plate were collected as one biological sample. Total RNA was extracted using the E.Z.N.A. Plant RNA kit with DNA digestion on column (Biotek, China) according to the manufacturer’s instructions. RNA samples were quantified with a Nanodrop spectrophotometer (ThermoFisher, USA). First strand cDNA was synthesized using the iScript^TM^ cDNA synthesis kit (Bio-Rad). qRT-PCR was performed using the iTaq^TM^ Universal SYBR Green Supermix (Bio-Rad) and CFX96 Real-time system (Bio-Rad) and the qPCR data was analyzed as previously described (Livak & Schmittgen, 2001; Wang et al, 2019). Primers for qPCR of SA-JA related genes in *N. benthamiana* were used as described by Nakano and Mukaihara (2018). Primer sequences are listed in Table S1.

### Measurements of SA and JA content in plant leaves

SA and JA content were quantified using the method described by Forcat and collaborators (2008) with the following modifications. Leaves (50 mg FW) were collected 42 hours after Agrobacterium infiltration and frozen in liquid nitrogen before grounding into fine powder with the Qiagen tissue lyser. SA and JA were extracted at 10。C for 1 h using 70% methanol extraction solvent spiked with d4-SA as internal standards. Supernatant was taken after centrifugation at 20000 rcf for 10 min and analyzed on ACQUITY UPLC I-class coupled with AB SCIEX TripleTOF 5600+. The analytical column used was a ACQUITY UPLC BECH C18 1.7 µm, 2.1X150 mm column. The JA concentration was calculated based on the calibration curve created by running a JA standard solution. The results were analyzed by Peakview1.2.

### Measurements of total cellular glutathione in *N. benthamiana* leaves

Total cellular glutathione was measured as previously described (Sang et al, 2016). Briefly, 10 mg of *N. benthamiana* leaves were collected and glutathione was measured using a Glutathione Assay Kit (Beyotime, China) according to the manufacturer’s instructions.

### Virus-induced gene silencing (VIGS) in *N. benthamiana*

VIGS in *N. benthamiana* plants was performed using TRV vectors as described (Senthil-Kumar & Mysore, 2014). VIGS of *NbSGT1* was performed with several modifications described by Yu and collaborators (2019). Cultures of Agrobacterium carrying pTRV2:*NbSGT1* plasmids or pTRV2 plasmids were mixed at 1:1 ratio and co-infiltrated into the lower leaves of 3-week-old *N. benthamiana* plants. The upper leaves were used for experimental assay within 7-10 days after VIGS application. Silencing of NRCs (NLR required for cell death) in *N. benthamiana* and subsequent expression of T3Es was performed as described by Wu and collaborators (2017).

### Pseudomonas syringae virulence assays

For leaf infiltration with *P. syringae*, Arabidopsis plants were treated with 100 µM EST for 2 days before inoculation. Plants showed no difference in root or shoot size at the time of inoculation. *Pto* DC3000 was resuspended in water at 10^5^ cfu/ml. The bacterial suspensions were then infiltrated into 4-to-5-week-old Arabidopsis leaves using a needleless syringe. For spray inoculation, *Pto* DC3000 was resuspended in water at 10^8^ cfu/ml, and silwet-L77 was added to a final concentration of 0.02% before spraying onto 3-week-old Arabidopsis seedlings. Bacterial numbers were determined 3 days post-inoculation as previously described (Macho *et al*., 2012; Wang *et al*., 2019).

### Ralstonia solanacearum virulence assays

For standard *R. solanacearum* virulence assays, 4-week-old *A. thaliana* plants, grown in Jiffy pots, were inoculated with *R. solanacearum* without wounding by soil drenching. For experiments using inducible transgenic lines, all the plants were treated with 100 µM EST for 2 days before inoculation. Plants showed no difference in root or shoot size at the time of inoculation. An overnight-grown bacterial suspension was diluted to obtain an inoculum of 5×10^7^ cfu/ml. Once the Jiffy pots were completely drenched, the plants were removed from the bacterial solution and placed back on a bed of potting mixture soil. The genotypes to be tested were placed in a random order in order to allow an unbiased analysis of the wilting. Daily scoring of the visible wilting on a scale ranging from 0 to 4 (or 0 to 100% leaves wilting) led to an analysis using the Kaplan-Meier survival analysis, log-rank test and hazard ratio calculation as previously described (Morel et al., 2018b).

To determine *R. solanacearum* growth in Arabidopsis leaves, a 10^7^ cfu/ml inoculum was infiltrated into leaves of 4-week-old Arabidopsis plants 2 days after EST treatment, and samples were taken 2 days after inoculation. To determine *R. solanacearum* growth in *N. benthamiana* leaves, a 10^5^ cfu/ml inoculum of *R. solanacearum* Y45 was infiltrated into *N. benthamiana* leaves expressing RipE1-GFP or a GFP control. RipE1-GFP was expressed using Agrobacterium, and *R. solanacearum* Y45 was infiltrated in leaf tissues 24 hours after Agrobacterium infiltration, before the development of cell death. *R. solanacearum* Y45 is a strain originally isolated from tobacco (Li *et al*., 2011), which is pathogenic in *N. benthamiana* (unpublished data). To determine bacterial numbers, leaf discs (3 leaf discs from Arabidopsis plants and 4 leaf discs from *N. benthamiana* plants) were taken and weighed. The plant tissue was ground and homogenized in distilled water before plating serial dilutions to determine cfu per gram of fresh weight.

## Acknowledgements

We thank Nemo Peeters and Anne-Claire Cazale for sharing unpublished biological materials, Longjiang Fan, Yong Liu, Chanhong Kim, Alex Schultink, and Brian Staskawicz for sharing biological materials, Rosa Lozano-Duran for critical reading of this manuscript, and Xinyu Jian for technical and administrative assistance during this work. We thank the PSC Cell Biology, Proteomics, and Metabolomics core facilities for assistance with confocal microscopy and mass spectrometry analysis, respectively. This work was supported by the Strategic Priority Research Program of the Chinese Academy of Sciences (grant XDB27040204), the National Natural Science Foundation of China (NSFC; grant 31571973), the Chinese 1000 Talents Program, and the Shanghai Center for Plant Stress Biology (Chinese Academy of Sciences). The authors have no conflict of interest to declare.

---

**Figure S1.**
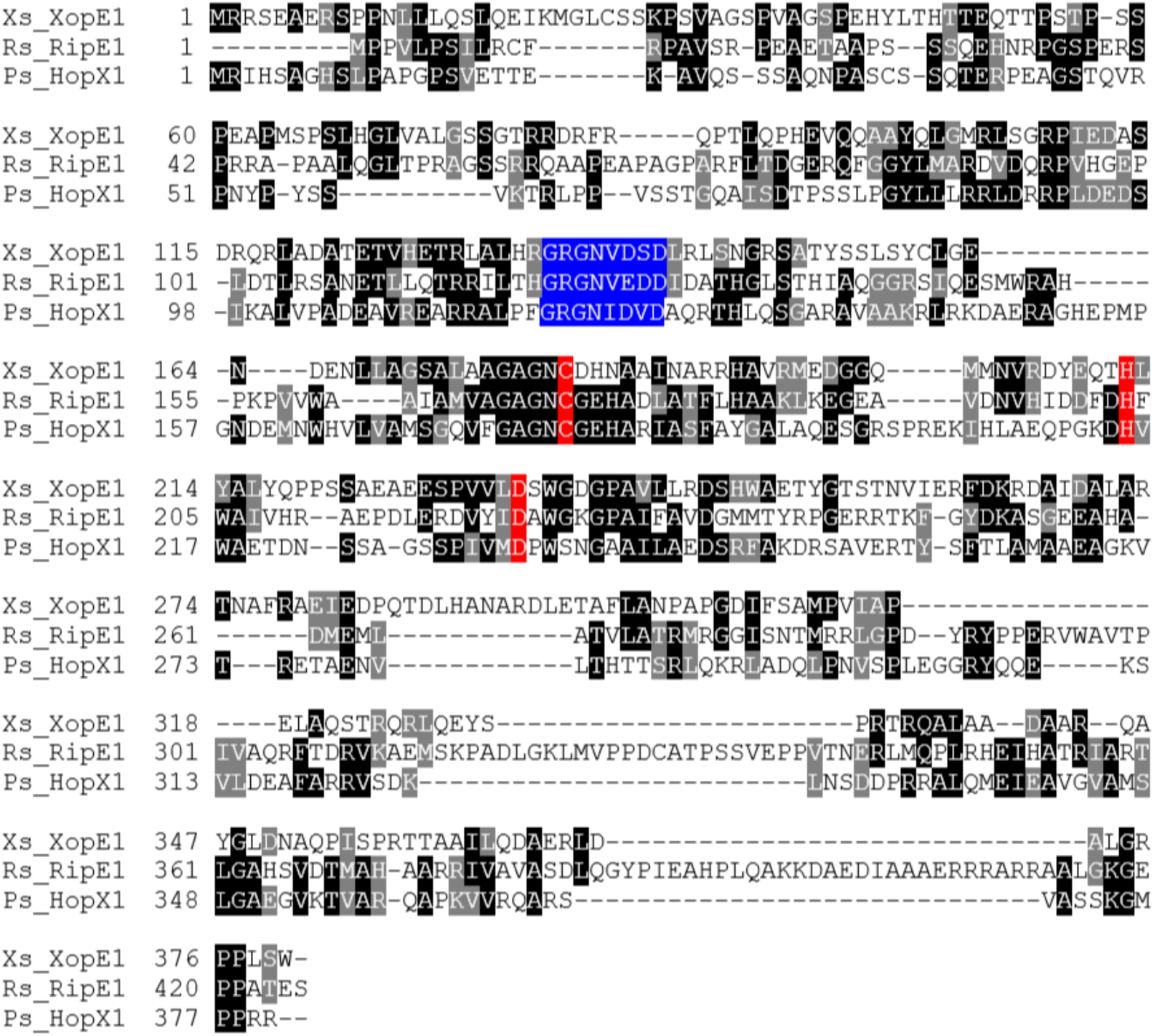
Phylogenetic analysis of RipE1. Alignment of the amino acid sequence of RipE1 from *R. solanacearum* GMI1000 (Rs_RipE1), XopE1 from *Xanthomonas campestris* pv. *vesicatoria* (Xs_XopE1), and HopX1 from *Pseudomonas syringae* pv tabaci 11528 (Ps_HopX1). Residues forming the predicted catalytic triad are indicated in red, and the conserved domain A is indicated in blue. The black shaded amino acids are identical among the three effectors.

**Figure S2.**
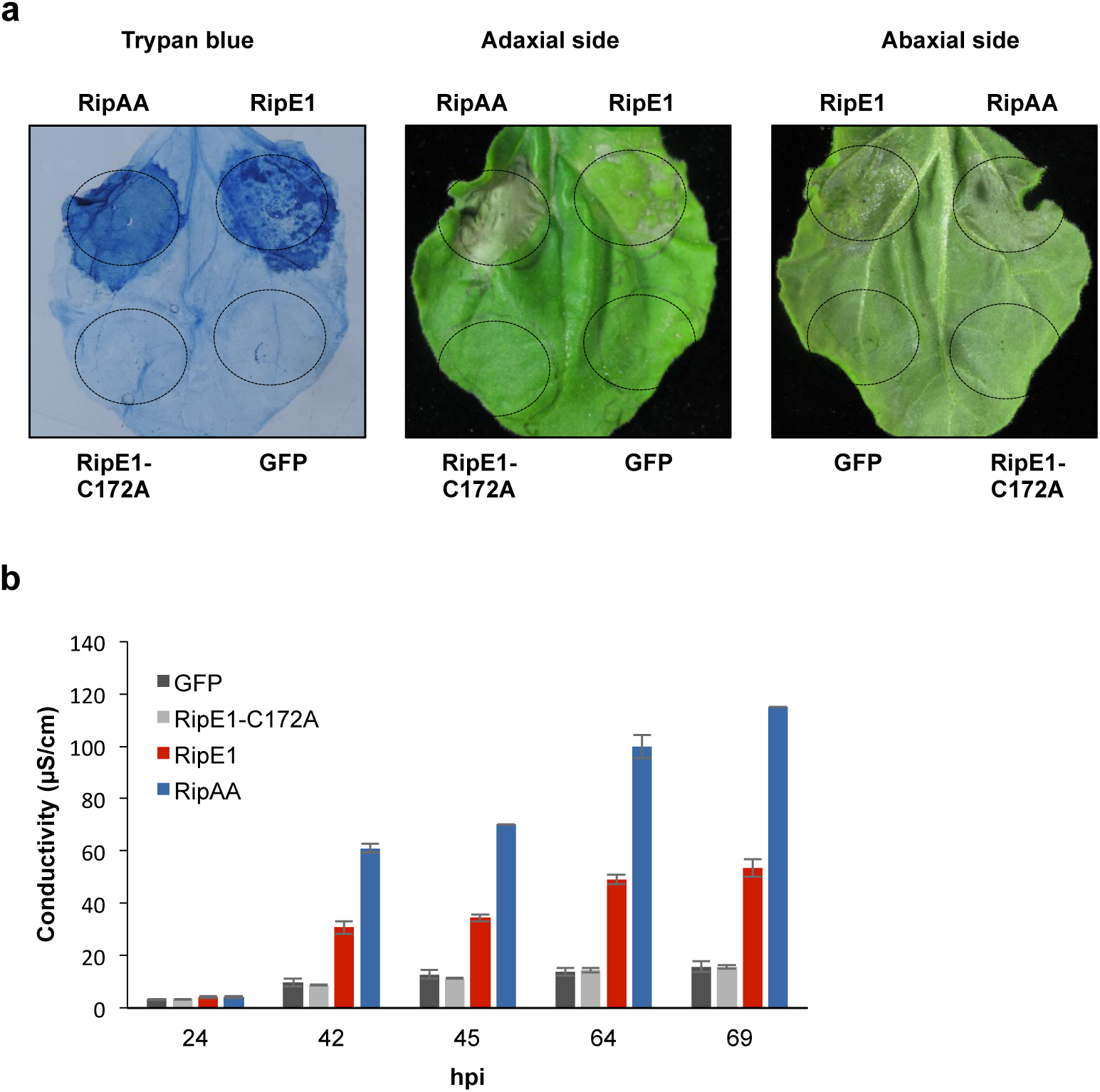
RipE1 induces cell death in *N. benthamiana.* RipE1-GFP, RipE1-C172A-GFP, RipAA-GFP, and GFP (as control) were expressed in the same leaf of *N. benthamiana* using Agrobacterium with an OD_600_ of 0.5. (a) Trypan blue staining was performed 4 days post-inoculation, and additional photos of an independent unstained leaf is shown for reference. (b) Ion leakage measured in leaf discs taken from *N. benthamiana* tissues expressing the indicated constructs, at the indicated time points. Values indicate mean ± SE (n=3 biological replicates). These experiments were repeated 3 times with similar results.

**Figure S3.**
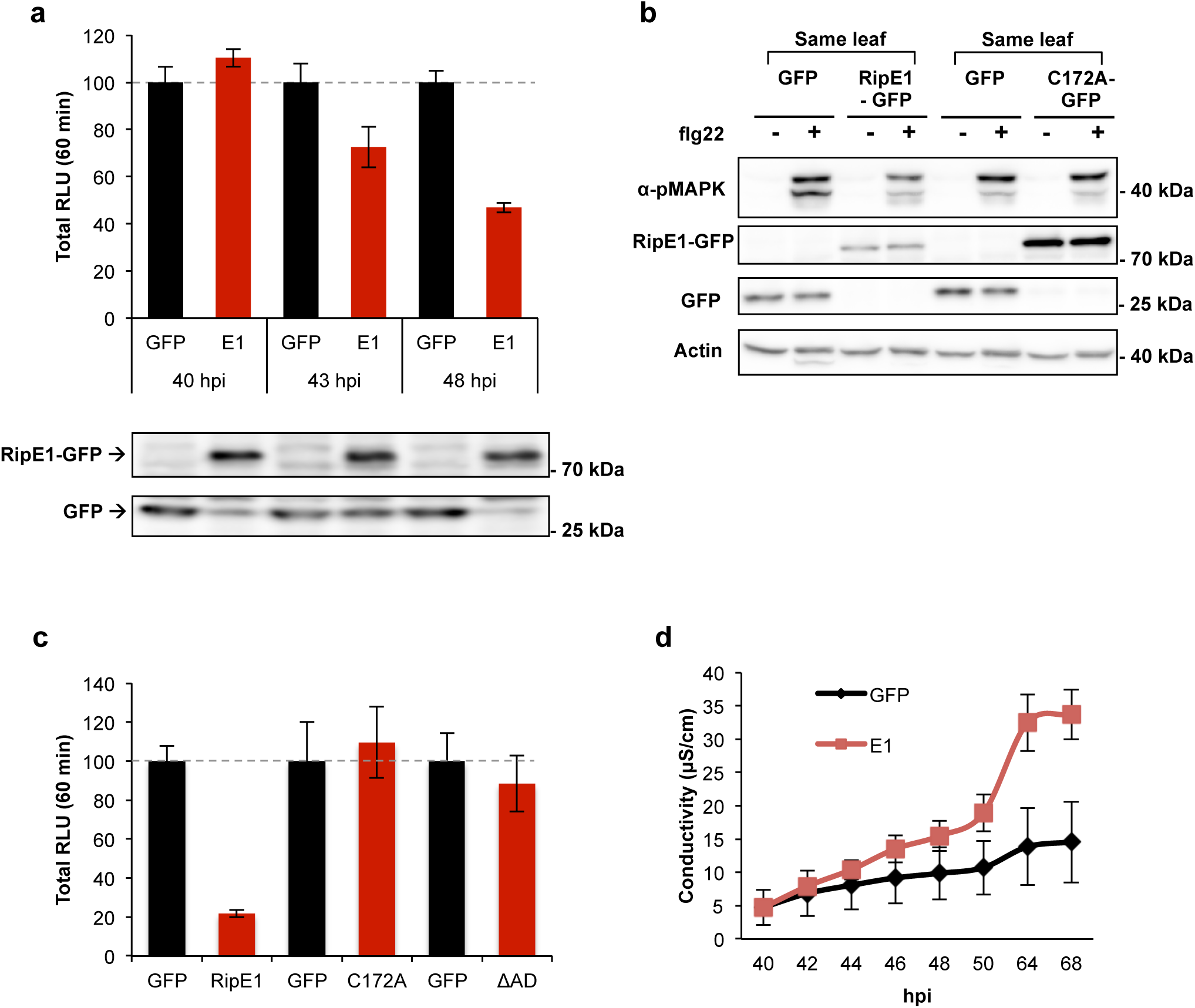
RipE1 expression inhibits PTI responses in *N. benthamiana*, which correlates with the induction of cell death. Agrobacterium was used to induce the transient expression of RipE1-GFP in half of the leaf and GFP in the other half. (a) Oxidative burst triggered by 50 nM flg22 in *N. benthamiana* tissues at the indicated time points, measured in a luminol-based assay as relative luminescence units (RLU). Values are average ± SE (n=24), and are represented as % of the GFP control in each time point. Western blot with anti-GFP is shown for reference of protein accumulation at each time point. (b) MAPK activation was induced 40 hours after Agrobacterium infiltration with 100 nM flg22 and analysed 15 minutes after flg22 treatment using anti-phosphorylated MAPK antibody (anti-pMAPK). Immunoblots were also analysed using anti-GFP antibody to verify protein accumulation. Anti-actin was used to verify equal loading. Molecular weight (kDa) marker bands are indicated for reference. (c) Oxidative burst was induced as in (a) and measured 2 days post-Agrobacterium infiltration. Mutant variants are described in the Figure 1. (d) Ion leakage was measured as in the Figure 1. Measurement over time after RipE1 expression reflects that the induction of cell death correlates in time with the suppression of PTI responses. The experiments were repeated three times with similar results.

**Figure S4.**
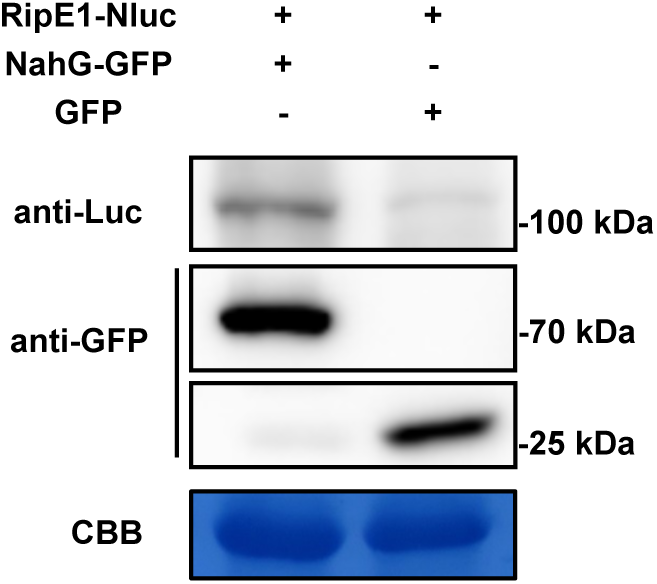
Protein accumulation upon co-expression of RipE1 and NahG. Western blot showing protein accumulation in the experiments shown in the figure 2. Molecular weight (kDa) marker bands are indicated for reference.

**Figure S5.**
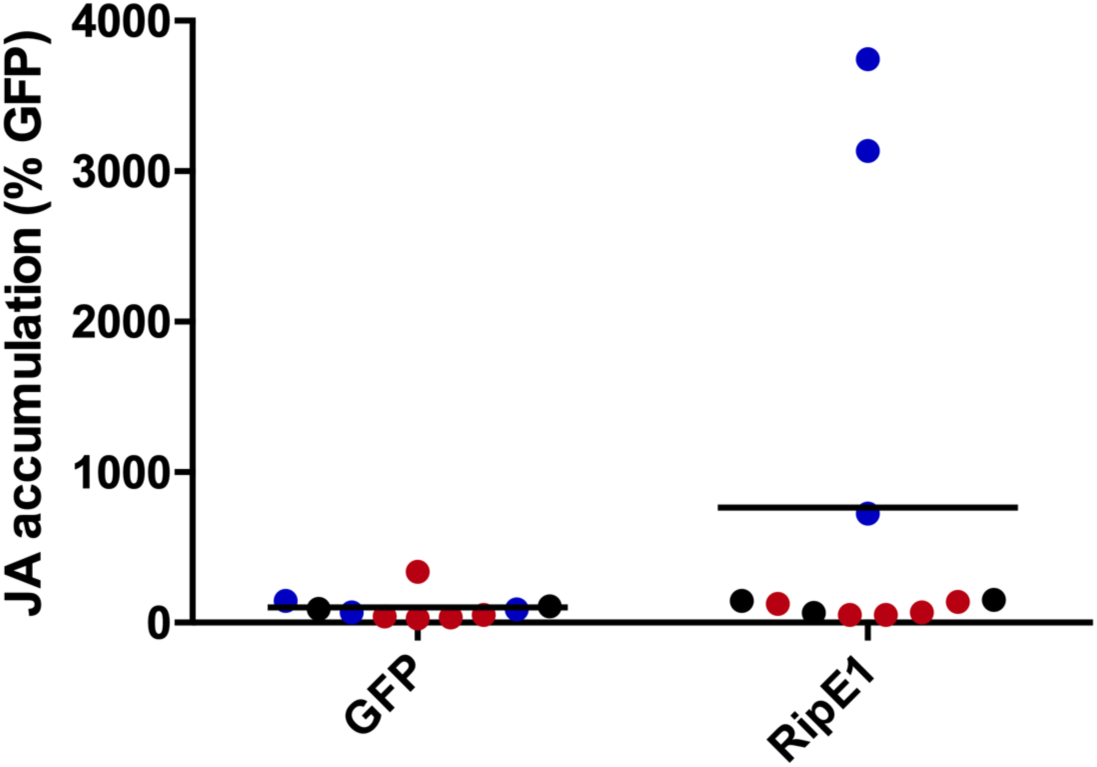
RipE1 expression leads to an increase in JA contents. Measurement of JA accumulation in *N. benthamiana* tissues expressing GFP or RipE1, using Agrobacterium with an OD_600_ of 0.5. Samples were taken 42 hours after Agrobacterium infiltration. Three independent biological repeats were performed, and the different colors indicate values from different replicates. Values are represented as % of the GFP control in each replicate.

**Figure S6.**
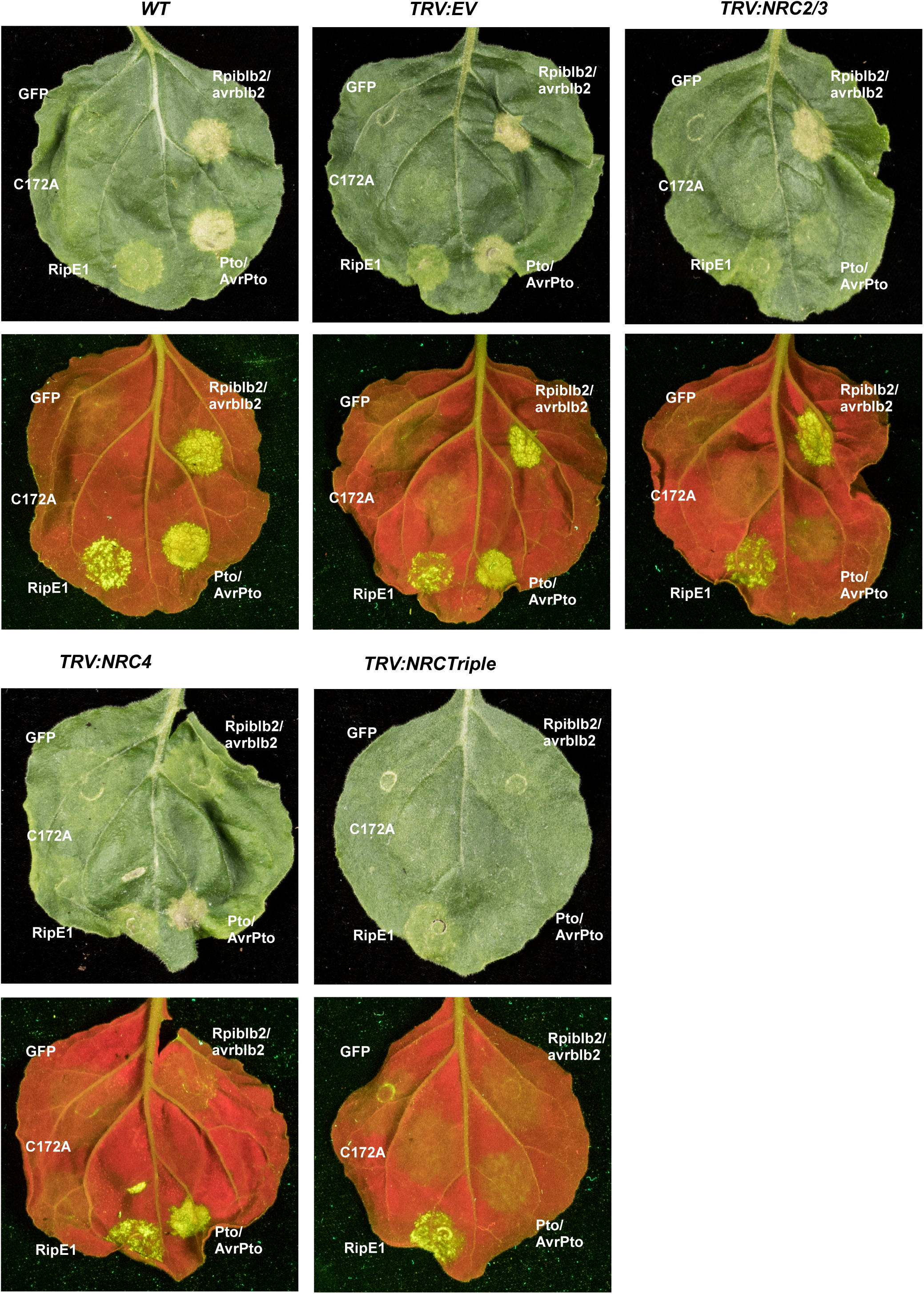
RipE1-triggered cell death does not require NRC proteins. RipE1-GFP, RipE1-C172A-GFP or GFP (as control) were transiently expressed using agrobacterium into wild type (WT) *N. benthamiana*, leaves silenced with EV (as control) and those silenced with different NRC homologs (NRC2/3, NRC4 and NRC2/3/4-Triple), using VIGS. For RipE1-GFP, RipE1-C172A-GFP and GFP an OD_600_ of 0.5 was used. Rpiblb2 (OD_600_ 0.2)/AVRblb2 (OD_600_ 0.1) and Pto (OD_600_ 0.6)/AVRPto (OD_600_ 0.1), which are NRC4 and NRC2/3 dependent, respectively, were included as controls. Photos were taken 5 days post inoculation under natural or UV light. UV images were taken from the abaxial side and flipped horizontally for representation.

**Figure S7.**
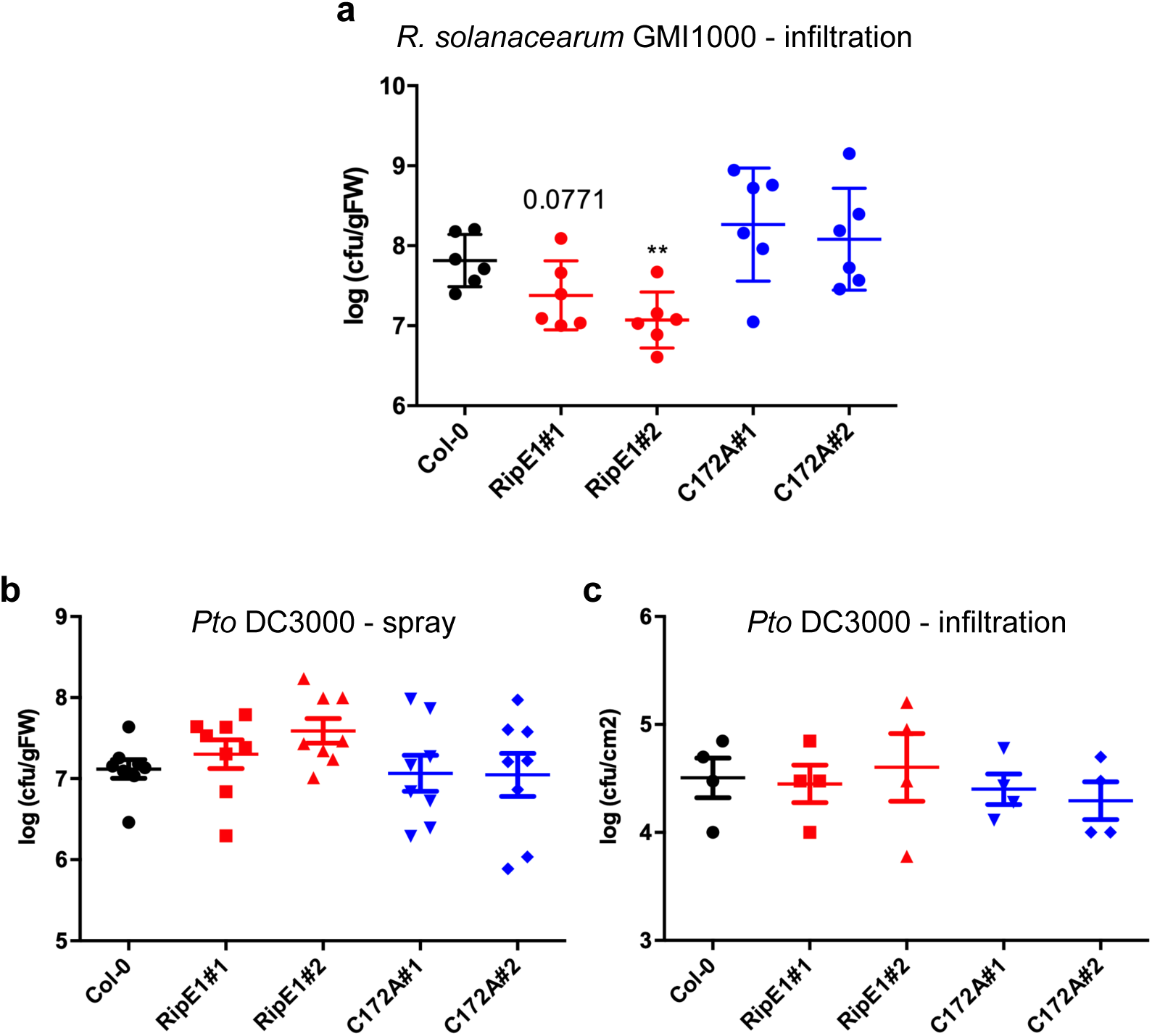
Leaves of RipE1-expressing Arabidopsis plants show enhanced resistance against inoculation with *R. solanacearum*, but not *P. syringae pv tomato* (*Pto*) DC3000. (a) Growth of *R. solanacearum* GMI1000 (inoculated at 10^7^ cfu/ml) infiltrated with a needleless syringe into wild-type Col-0 and RipE1-expressing Arabidopsis plants. Four-week-old plants were sprayed with 100 µM estradiol and then covered for 2 days prior inoculation. Bacterial colony-forming units (cfu) were determined 2 days post-inoculation (dpi). Values indicate mean ± SE (n=6 biological replicates). Asterisks indicate significant differences compared to Col-0 WT plants according to a Student’s t test (** p < 0.01). A specific p value is indicated for the RipE1#1 plant to indicate a reproducible (but not statistically significant) difference in bacterial growth compared to Col-0 WT plants. (b) Growth of surface (spray)-inoculated *Pto* DC3000 (inoculated at 10^8^ cfu/ml) in wild-type Col-0 and RipE1-expressing Arabidopsis seedlings. Three-week-old seedlings were sprayed with 100 µM estradiol and then covered for 2 days prior inoculation. Bacterial colony-forming units (cfu) were determined 3 days post-inoculation (dpi). Values indicate mean ± SE (n=8 biological replicates). (c) Growth of *Pto* DC3000 (inoculated at 10^5^ cfu/ml) infiltrated with a needleless syringe into wild-type Col-0 and RipE1-expressing Arabidopsis plants. Four-week-old plants were sprayed with 100 µM estradiol and then covered for 2 days prior inoculation. Bacterial colony-forming units (cfu) were determined 3 days post-inoculation (dpi). Values indicate mean ± SE (n=4 biological replicates). Experiments were repeated more than three times with similar results.

**Figure S8.**
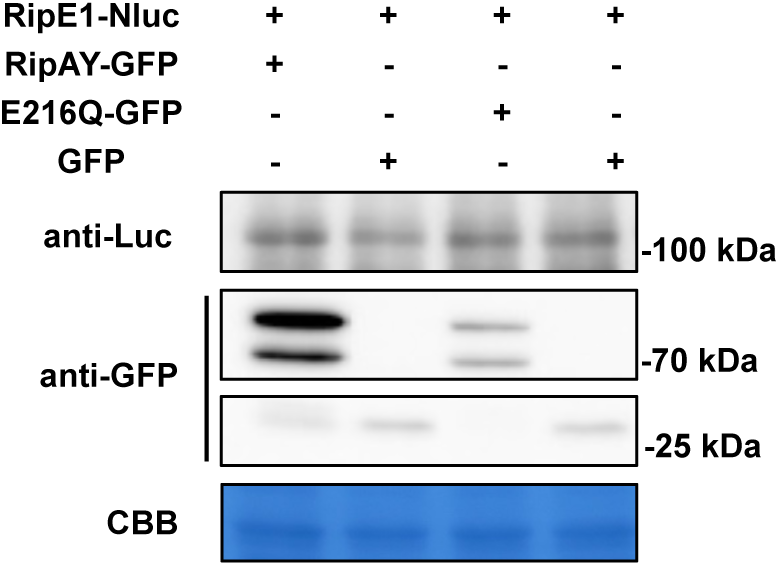
Protein accumulation upon co-expression of RipE1 and RipAY. Western blot showing protein accumulation in the experiments shown in the figure 6. Molecular weight (kDa) marker bands are indicated for reference.

**Figure S9.**
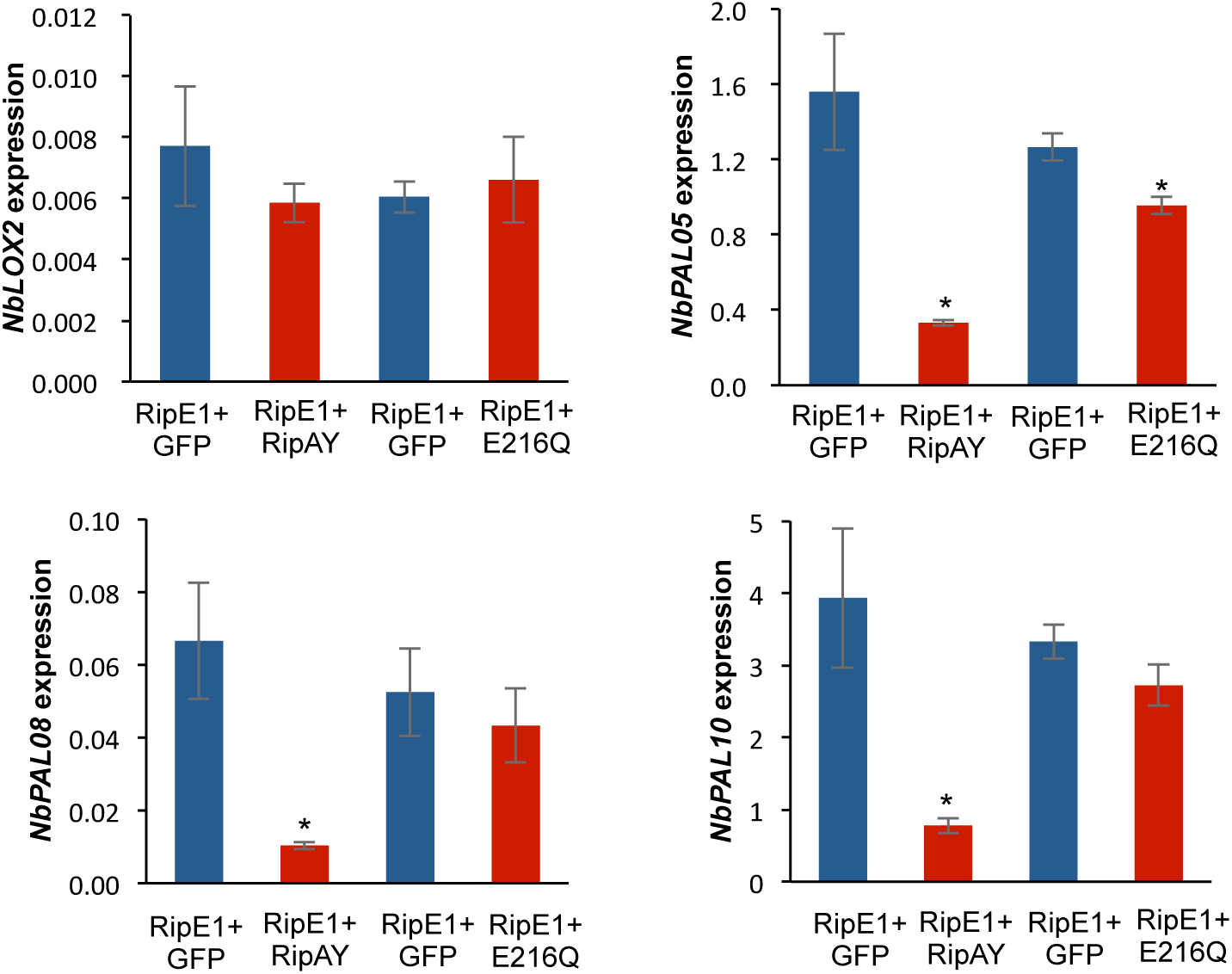
RipAY suppresses the overexpression of SA-related genes triggered by RipE1. Quantitative RT-PCR to determine the expression of *NbPAL05, NbPAL08, NbPAL10,* and *NbLOX2* in *N. benthamiana* tissues 48 hours after Agrobacterium infiltration. *NbPAL05:* Niben101Scf05617g00005.1; *NbPAL08*: Niben101Scf03712g01008.1; *NbPAL10:* Niben101Scf12881g00010.1. Expression values are relative to the expression of the housekeeping gene *NbEF1a*. Values indicate mean ± SE (n=3 biological replicates). Asterisks indicate significant differences compared to the mock control according to a Student’s t test (* p < 0.05).

**Table S1:**
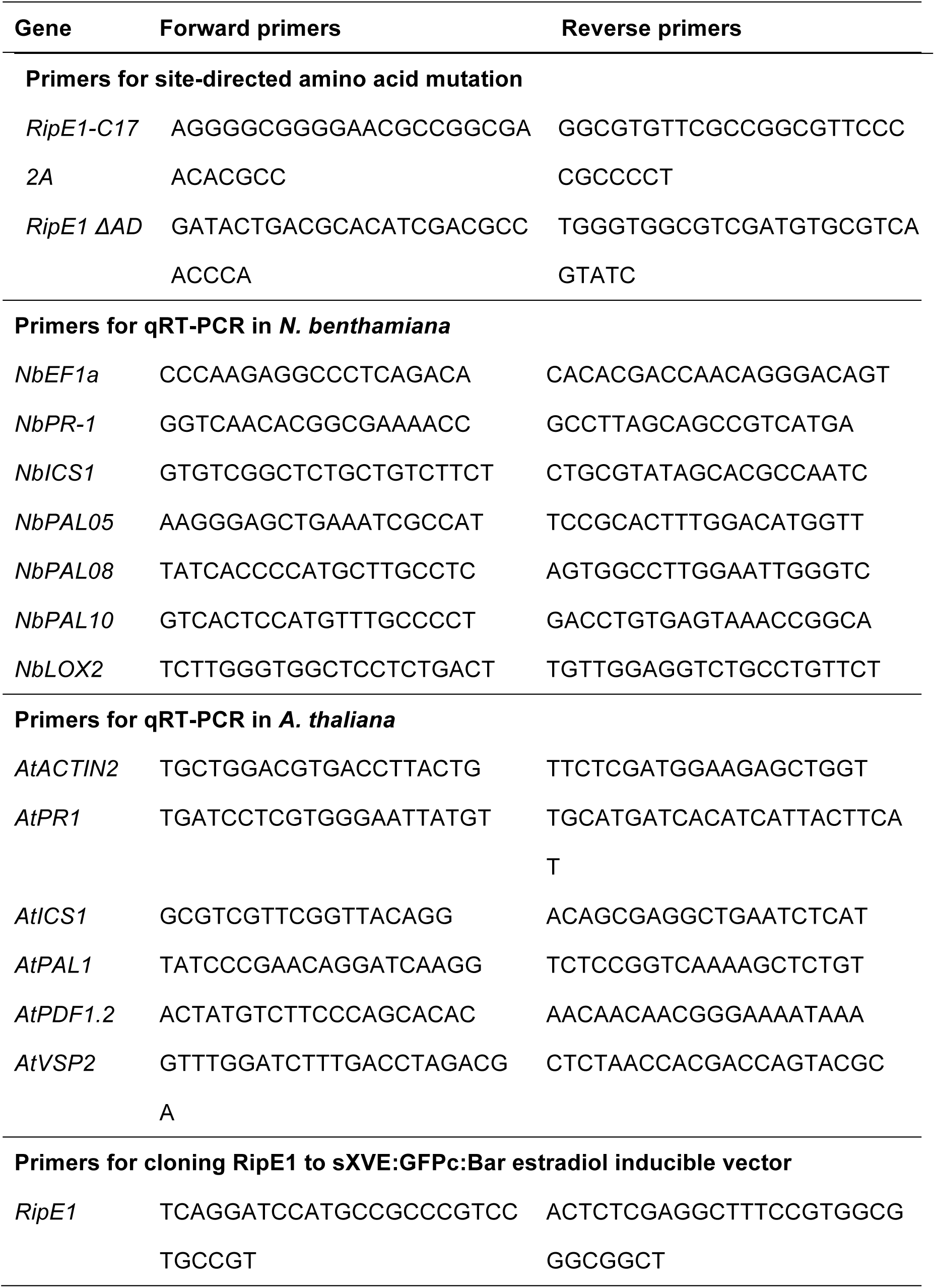
Primers used in this study for RipE1 cloning and qRT-PCR.

## References

1. Azevedo, C., Sadanandom, A., Kitagawa, K., Freialdenhoven, A., Shirasu, K., and Schulze-Lefert, P. (2002). The RAR1 interactor SGT1, an essential component of *R* gene-triggered disease resistance. Science 295:2073–2076.

2. Bell E, Creelman RA, Mullet JE. (1995). A chloroplast lipoxygenase is required for wound-induced jasmonic acid accumulation in Arabidopsis. Proceedings of the National Academy of Sciences, USA 92: 8675–8679.

3. Burger, M., and Chory, J. (2019). Stressed out about hormones: how plants orchestrate immunity. Cell Host Microbe 26:163–172.

4. Catherine, D., Yves, M., Nicolas, D., Marine, C., Patrick, D., Philippe, R., Yves, M., Alain, J., and Deborah, G. (2012). Deciphering the route of *Ralstonia solanacearum* colonization in *Arabidopsis thaliana* roots during a compatible interaction: focus at the plant cell wall. Planta 236:1419–1431.

5. Chiang, Y.-H., and Coaker, G. (2015). Effector Triggered Immunity: NLR immune perception and downstream defense responses. The Arabidopsis Book 2015.

6. Clarke, C.R., Studholme, D.J., Byron, H., Brendan, R., Alexandra, W., Rongman, C., Tadeusz, W., Marie-Christine, D., Emmanuel, W., and Castillo, J.A. (2015). Genome-enabled phylogeographic investigation of the quarantine pathogen *Ralstonia solanacearum* Race 3 Biovar 2 and screening for sources of resistance against its core effectors. Phytopathology 105:597–607.

7. Cui, H., Tsuda, K., Parker, J.E. (2015). Effector-triggered immunity: from pathogen perception to robust defense. In: Annual Review Plant Biology. 487–511.

8. Delaney, T.P., Uknes, S., Vernooij, B., Friedrich, L., Weymann, K., Negrotto, D., Gaffney, T., Gut-Rella, M., Kessmann, H., and Ward, E. (1994). A central role of salicylic acid in plant disease resistance. Science 266:1247–1250.

9. Forcat, S., Bennett, M., Mansfield, J., and Grant, M. (2008). A rapid and robust method for simultaneously measuring changes in the phytohormones ABA, JA and SA in plants following biotic and abiotic stress. Plant methods 4:16.

10. Fujiwara, S., Kawazoe, T., Ohnishi, K., Kitagawa, T., Popa, C., Valls, M., Genin, S., Nakamura, K., Kuramitsu, Y., and Tanaka, N. (2016). RipAY, a plant pathogen effector protein exhibits robust γ-glutamyl cyclotransferase activity when stimulated by eukaryotic thioredoxins. Journal of Biological Chemistry 291:6813–6830.

11. Galán, J.E., Lara-Tejero, M., Marlovits, T.C., and Wagner, S. (2014). Bacterial type III secretion systems: specialized nanomachines for protein delivery into target cells. Annual Review of Microbiology 68:415–438.

12. Garcion, C., Lohmann, A., Lamodière, E., Catinot, J., Buchala, A., Doermann, P., and Métraux, J.-P. (2008). Characterization and biological function of the ISOCHORISMATE SYNTHASE2 gene of Arabidopsis. Plant Physiology 147:1279–1287.

13. Gassmann, W., ., Hinsch, M.E., and Staskawicz, B.J. (2010). The Arabidopsis RPS4 bacterial-resistance gene is a member of the TIR-NBS-LRR family of disease-resistance genes. Plant Journal for Cell & Molecular Biology 20:265–277.

14. Gimenez-Ibanez, S., Boter, M., Fernández-Barbero, G., Chini, A., Rathjen, J.P., and Solano, R. (2014). The bacterial effector HopX1 targets JAZ transcriptional repressors to activate jasmonate signaling and promote infection in Arabidopsis. PLoS Biology 12:e1001792–e1001792.

15. Gimenez-Ibanez, S., Chini, A., & Solano, R. (2016). How microbes twist jasmonate signaling around their little fingers. Plants (Basel, Switzerland), 5: 9.

16. Jayaraman, J., Segonzac, C., Cho, H., Jung, G., and Sohn, K.H. (2016). Effector-assisted breeding for bacterial wilt resistance in horticultural crops. Horticulture Environment & Biotechnology 57:415–423.

17. Jiang, G., Wei, Z., Xu, J., Chen, H., Zhang, Y., She, X., Macho, A.P., Ding, W., and Liao, B. (2017). Bacterial wilt in China: history, current status, and future perspectives. Frontiers in Plant Science 8:1549.

18. Jones, J.D.G., and Dangl, J.L. (2006). The plant immune system. Nature 444:323–329.

19. Kadota, Y., Shirasu, K., and Guerois, R. (2010). NLR sensors meet at the SGT1– HSP90 crossroad. Trends in biochemical sciences 35:199–207.

20. Lavie, M., Shillington, E., Eguiluz, C., Grimsley, N., and Boucher, C. (2002). PopP1, a new member of the YopJ/AvrRxv family of type III effector proteins, acts as a host-specificity factor and modulates aggressiveness of *Ralstonia solanacearum*. Molecular Plant Microbe Interactions :15:1058–1068.

21. Leroux, C., Huet, G., Jauneau, A., Camborde, L., Trémousaygue, D., Kraut, A., Zhou, B., Levaillant, M., Adachi, H., and Yoshioka, H. (2015). A receptor pair with an integrated decoy converts pathogen disabling of transcription factors to immunity. Cell 161:1074–1088.

22. Li Z, Wu S, Bai X, Liu Y, Lu J, Liu Y, Xiao B, Lu X, Fan L. (2011) Genome sequence of the tobacco bacterial wilt pathogen Ralstonia solanacearum. Journal of Bacteriology 193: 6088–6089.

23. Li, X. (2011). Infiltration of Nicotiana benthamiana protocol for transient expression via Agrobacterium. Bio-protocol Bio101:e95.

24. Livak, K.J., and Schmittgen, T.D. (2001). Analysis of relative gene expression data using real-time quantitative PCR and the 2(-Delta Delta C(T)) Method. Methods 25:402–408.

25. Lv, R., Li, Z., Li, M., Dogra, V., Lv, S., Liu, R., Lee, K.P., and Kim, C. (2019) Uncoupled expression of nuclear and plastid photosynthesis-associated genes contributes to cell death in a lesion mimic mutant. Plant Cell 31: 210– 230.

26. Macho, A.P., Boutrot, F., Rathjen, J.P., and Zipfel, C. (2012). ASPARTATE OXIDASE plays an important role in Arabidopsis stomatal immunity. Plant Physiology 159: 1845–1856.

27. Macho, A.P. (2016). Subversion of plant cellular functions by bacterial type-III effectors: beyond suppression of immunity. New Phytologist 210:51–57.

28. Macho, A.P., and Zipfel, C. (2015). Targeting of plant pattern recognition receptor-triggered immunity by bacterial type-III secretion system effectors. Current Opinion in Microbiology 23:14–22.

29. Makarova, K.S., Aravind, L., and Koonin, E.V. (1999). A superfamily of archaeal, bacterial, and eukaryotic proteins homologous to animal transglutaminases. Protein Science 8:1714–1719.

30. Manifield, John, Genin, Stephane, Magori, Shimpei, Citovsky, Vitaly, and Sriariyanum. (2012). Top 10 plant pathogenic bacteria in molecular plant pathology. Molecular Plant Pathology 13:614–629.

31. Mansfield, J., Jenner, C., Hockenhull, R., Bennett, M.A., and Stewart, R. (1994). Characterization of avrPphE, a gene for cultivar-specific avirulence from *Pseudomonas syringae* pv. *phaseolicola* which is physically linked to hrpY, a new hrp gene identified in the halo-blight bacterium. Molecular Plant Microbe Interaction 7:726–739.

32. Morel A, G.J., Lonjon F, Sujeeun L, Barberis P, Genin S, Vailleau F, Daunay MC, Dintinger J, Poussier S, Peeters N, Wicker E. (2018a). The eggplant AG91-25 recognizes the Type III-secreted effector RipAX2 to trigger resistance to bacterial wilt (*Ralstonia solanacearum* species complex). Molecular Plant Pathology 19:2459–2472.

33. Morel, A., Peeters, N., Vailleau, F., Barberis, P., Jiang, G., Berthomé, R., and Guidot, A. (2018b). Plant pathogenicity phenotyping of *Ralstonia solanacearum* strains. in: Host-Pathogen Interactions: Methods and Protocols--Medina, C., and López-Baena, F.J., eds. New York, NY: Springer New York. 223–239.

34. Mukaihara, T., Hatanaka, T., Nakano, M., and Oda, K. (2016). *Ralstonia solanacearum* Type III effector RipAY is a glutathione-degrading enzyme that is activated by plant cytosolic thioredoxins and suppresses plant immunity. Mbio 7:e00359.

35. Mukaihara, T., Tamura, N., and Iwabuchi, M. (2010). Genome-wide identification of a large repertoire of *Ralstonia solanacearum* type III effector proteins by a new functional screen. Molecular Plant Microbe Interaction 23:251–262.

36. Nahar, K., Matsumoto, I., Taguchi, F., Inagaki, Y., Yamamoto, M., Toyoda, K., Shiraishi, T., Ichinose, Y., and Mukaihara, T. (2014). *Ralstonia solanacearum* type III secretion system effector Rip36 induces a hypersensitive response in the nonhost wild eggplant *Solanum torvum*. Molecular Plant Pathology 15:297–303.

37. Nakagawa, T., Suzuki, T., Murata, S., Nakamura, S., Hino, T., Maeo, K., Tabata, R., Kawai, T., Tanaka, K., and Niwa, Y. (2007). Improved Gateway binary vectors: high-performance vectors for creation of fusion constructs in transgenic analysis of plants. Journal of the Agricultural Chemical Society of Japan 71:2095–2100.

38. Nakano M, Mukaihara T. (2019). The type III effector RipB from *Ralstonia solanacearum* RS1000 acts as a major avirulence factor in *Nicotiana benthamiana* and other Nicotiana species. Molecular Plant Pathology 20:1237–1251.

39. Nakano, M., and Mukaihara, T. (2018). *Ralstonia solanacearum* Type III effector RipAL targets chloroplasts and induces jasmonic acid production to suppress salicylic acid-mediated defense responses in plants. Plant and Cell Physiology 59:2576–2589.

40. Nimchuk, Z.L., Fisher, E.J., Desveaux, D., Chang, J.H., and Dangl, J.L. (2007). The HopX (AvrPphE) family of *Pseudomonas syringae* type III effectors require a catalytic triad and a novel N-terminal domain for function. Molecular Plant Microbe Interactions 20:346–357.

41. Peeters, N., Carrère, S., Anisimova, M., Plener, L., Cazalé, A.C., and Genin, S. (2013a). Repertoire, unified nomenclature and evolution of the Type III effector gene set in the *Ralstonia solanacearum* species complex. BMC Genomics 14:859–859.

42. Peeters, N., Guidot, A., Vailleau, F., and Valls, M. (2013b). *Ralstonia solanacearum*, a widespread bacterial plant pathogen in the post-genomic era. Molecular Plant Pathology 14:651–662.

43. Poueymiro, M., Cunnac, S., Barberis, P., Deslandes, L., Peeters, N., Cazale-Noel, A.C., Boucher, C., and Genin, S. (2009). Two type III secretion system effectors from *Ralstonia solanacearum* GMI1000 determine host-range specificity on tobacco. Molecular Plant Microbe Interaction 22:538–550.

44. Rosas-Díaz, T., Cana-Quijada, P., Amorim-Silva, V., Botella, M.A., Lozano-Durán, R., and Bejarano, E.R. (2017). *Arabidopsis NahG* plants as a suitable and efficient system for transient expression using *Agrobacterium tumefaciens*. Molecular Plant 10:353–356.

45. Rufián, J.S, Lucía, A, Rueda-Blanco, J, Zumaquero, A, Guevara, C.M, Ortiz-Martín, I, Ruiz-Aldea, G, Macho, A.P, Beuzón, C.R, and Ruiz-Albert J (2018) Suppression of HopZ effector-triggered plant immunity in a natural pathosystem. Frontiers in Plant Sciences 9:977–981.

46. Sang, Y., and Macho, A.P. (2017). Analysis of PAMP-triggered ROS burst in plant immunity. Methods in Molecular Biology 1578:143–153.

47. Sang, Y., Wang, Y., Ni, H., Cazalã, A.C., She, Y.M., Peeters, N., and Macho, A.P. (2016). The *Ralstonia solanacearum* type III effector RipAY targets plant redox regulators to suppress immune responses. Molecular Plant Pathology 19:129–142.

48. Sarris, P., Duxbury, Z., Huh, S.U., Ma, Y., Segonzac, C., Sklenar, J., Derbyshire, P., Cevik, V., Rallapalli, G., and Saucet, S. (2015). A plant immune receptor detects pathogen effectors that target WRKY transcription factors. Cell 161:1089–1100.

49. Schlücking, K., Kai, H.E., Köster, P., Drerup, M.M., Eckert, C., Steinhorst, L., Waadt, R., Batistič, O., and Kudla, J. (2013). A new β-estradiol-inducible vector set that facilitates easy construction and efficient expression of transgenes reveals CBL3-dependent cytoplasm to tonoplast translocation of CIPK5. Molecular Plant 6:1814–1829.

50. Schultink, A., Qi, T., Lee, A., Steinbrenner, A.D., and Staskawicz, B. (2017). Roq1 mediates recognition of the Xanthomonas and Pseudomonas effector proteins XopQ and HopQ1. Plant Journal 92:787–795

51. Segonzac, C., Feike, D., Gimenez Ibanez, S., Dagmar R, H., Zipfel, C., and John P, R. (2011). Hierarchy and roles of pathogen-associated molecular pattern-induced responses in Nicotiana benthamiana. Plant Physiology 156:687–699.

52. Senthil-Kumar, M., and Mysore, K.S. (2014). Tobacco rattle virus–based virus-induced gene silencing in Nicotiana benthamiana. Nature Protocols 9:1549–1562.

53. Shine, M.B., Yang, J.W., El-Habbak, M., Nagyabhyru, P., Fu, D.Q., Navarre, D., Ghabrial, S., Kachroo, P., and Kachroo, A. (2016). Cooperative functioning between phenylalanine ammonia lyase and isochorismate synthase activities contributes to salicylic acid biosynthesis in soybean. New Phytologist 212:627–636.

54. Solé, M., Popa, C., Mith, O., Sohn, K.H., Jones, J., Deslandes, L., and Valls, M. (2012). The awr gene family encodes a novel class of *Ralstonia solanacearum* type III effectors displaying vrulence and avirulence activities. Molecular Plant Microbe Interactions 25:941–953.

55. Stevens, C., Bennett, M.A., Athanassopoulos, E., Tsiamis, G., Taylor, J.D., and Mansfield, J.W. (1998). Sequence variations in alleles of the avirulence gene avrPphE.R2 from *Pseudomonas syringae* pv. *phaseolicola* lead to loss of recognition of the AvrPphE protein within bean cells and a gain in cultivar-specific virulence. Molecular Microbiology 29:165–177.

56. Tasset, C., Bernoux, M., Jauneau, A., Pouzet, C., Brière, C., Kieffer-Jacquinod, S., Rivas, S., Marco, Y., and Deslandes, L. (2010). Autoacetylation of the *Ralstonia solanacearum* effector PopP2 targets a lysine residue essential for RRS1-R-mediated immunity in Arabidopsis. PLOS Pathogens 6:e1001202.

57. Turner, M., Jauneau, A., Genin, S., Tavella, M.J., Vailleau, F., Gentzbittel, L., and Jardinaud, M.F. (2009). Dissection of bacterial wilt on *Medicago truncatula* revealed two type III secretion system effectors acting on root infection process and disease development. Plant Physiology 150:1713–1722.

58. Vlot, A.C., Dempsey, D.M.A., and Klessig, D.F. (2009). Salicylic acid, a multifaceted hormone to combat disease. Annual Review of Phytopathology 47:177–206.

59. Wang, Y., Li, Y., Rosas-Diaz, T., Caceres-Moreno, C., Lozano-Durán, R., and Macho, A.P. (2019). The IMMUNE-ASSOCIATED NUCLEOTIDE-BINDING 9 protein is a regulator of basal immunity in *Arabidopsis thaliana*. Molecular Plant Microbe Interactions 32:65–75.

60. Ward, E.R., Uknes, S.J., Williams, S.C., Dincher, S.S., Wiederhold, D.L., Alexander, D.C., Ahlgoy, P., Métraux, J.P., and Ryals, J.A. (1991). Coordinate gene activity in response to agents that induce systemic acquired resistance. Plant Cell 3:1085–1094.

61. Wei, Y., Caceres-Moreno, C., Jimenez-Gongora, T., Wang, K., Sang, Y., Lozano-Duran, R., and Macho, A.P. (2017). The *Ralstonia solanacearum* csp22 peptide, but not flagellin-derived peptides, is perceived by plants from the Solanaceae family. Plant Biotechnology Journal 16:1349–1362

62. Wiermer, Marcel, Feys, Bart, J., Parker, and Jane, E. (2005). Plant immunity : the EDS1 regulatory node. Current Opinion in Plant Biology 8:383–389.

63. Wildermuth, M.C., Dewdney, J., ., Wu, G., ., and Ausubel, F.M. (2001). Isochorismate synthase is required to synthesize salicylic acid for plant defence. Nature 414:562–565.

64. Williams, S.J., Kee Hoon, S., Li, W., Maud, B., Sarris, P.F., Cecile, S., Thomas, V., Yan, M., Saucet, S.B., and Ericsson, D.J. (2014). Structural basis for assembly and function of a heterodimeric plant immune receptor. Science 344:299–303.

65. Wu, C.-H., Abd-El-Haliem, A., Bozkurt, T.O., Belhaj, K., Terauchi, R., Vossen, J.H., and Kamoun, S. (2017). NLR network mediates immunity to diverse plant pathogens. Proceedings of the National Academy of Sciences USA 114:8113–8118.

66. Wu, C.H., Belhaj, K., Bozkurt, T.O., Birk, M.S., and Kamoun, S. (2016). Helper NLR proteins NRC2a/b and NRC3 but not NRC1 are required for Pto-mediated cell death and resistance in *Nicotiana benthamiana*. New Phytologist 209:1344–1352.

67. Yu, G., Xian, L., Sang, Y., and Macho, A.P. (2019a). Cautionary notes on the use of Agrobacterium-mediated transient gene expression upon SGT1 silencing in *Nicotiana benthamiana*. New Phytologist 222:14–17.

68. Yu, G., Xian, L., Xue, H., Yu, W., Rufian, J., Sang, Y., Morcillo, R., Wang, Y., and Macho, A.P. (2019b). A bacterial effector protein prevents MAPK-mediated phosphorylation of SGT1 to suppress plant immunity. bioRxiv:641241.

69. Zhang, X., Henriques, R., Lin, S.-S., Niu, Q.-W., and Chua, N.-H. (2006). Agrobacterium-mediated transformation of *Arabidopsis thaliana* using the floral dip method. Nature Protocols 1:641–646.

